# Multivalent interactions with RNA drive RNA binding protein recruitment and dynamics in biomolecular condensates in *Xenopus* oocytes

**DOI:** 10.1101/2021.06.21.449303

**Authors:** Sarah E. Cabral, Kimberly L. Mowry

## Abstract

RNA localization and biomolecular condensate formation are key biological strategies for organizing the cytoplasm and generating cellular and developmental polarity. While enrichment of RNAs and RNA-binding proteins (RBPs) is a hallmark of both processes, the functional and structural roles of RNA-RNA and RNA-protein interactions within condensates remain unclear. Recent work from our laboratory has shown that RNAs required for germ layer patterning in *Xenopus* oocytes localize in novel biomolecular condensates, termed Localization bodies (L-bodies). L-bodies are composed of a non-dynamic RNA phase enmeshed in a more dynamic protein-containing phase. However, the interactions that drive the biophysical characteristics of L-bodies are not known. Here, we test the role of RNA-protein interactions using an L-body RNA-binding protein, PTBP3, which contains four RNA-binding domains (RBDs). We find that binding of RNA to PTB is required for both RNA and PTBP3 to be enriched in L-bodies *in vivo*. Importantly, while RNA binding to a single RBD is sufficient to drive PTBP3 localization to L-bodies, interactions between multiple RRMs and RNA tunes the dynamics of PTBP3 within L-bodies. *In vitro*, recombinant PTBP3 phase separates into non-dynamic structures in an RNA-dependent manner, supporting a role for RNA-protein interactions as a driver of both recruitment of components to L-bodies and the dynamics of the components after enrichment. Our results point to a model where RNA serves as a concentration-dependent, non-dynamic substructure and multivalent interactions with RNA are a key driver of protein dynamics.

## Introduction

Subcellular compartmentalization is a conserved mechanism by which cells enrich biomolecules for particular processes, allowing for spatial control of biological activity. Many of these compartments, including stress granules, nucleoli, and germ granules, are biomolecular condensates, enriching proteins and RNAs relative to their surroundings without a lipid membrane (reviewed in Boeynaems et al., 2018). The formation of biomolecular condensates is thought to be driven by multivalent interactions, either between “sticker” domains in protein intrinsically disordered regions (IDRs) or between ordered interaction domains, such as multivalent signaling or RNA-binding proteins (RBPs) (reviewed in Mittag and Parker, 2018). These interactions lead to biomolecular condensates with a wide range of biophysical states, with varying dynamics from liquid to gel to solid (reviewed in Alberti et al., 2019). However, the relative contributions of each of these interaction domains to the formation and dynamics of different types of condensates *in vivo* is unclear.

In addition to being enriched for certain types of proteins, a conserved feature of many classes of biomolecular condensates is incorporation of RNA (reviewed in Fay and Anderson, 2018). Emerging research, particularly *in vitro* studies, suggests that RNA—through both RNA-RNA and RNA-RBP intermolecular interactions—may play a critical role in the structure and assembly of biomolecular condensates (reviewed in Van Treeck and Parker, 2018). However, the role of RNAs and multivalent RBPs in *in vivo* condensates remains unclear and may vary based on the concentration of the RNA and type of condensate. *In vitro*, low concentrations of RNA often promote liquid-liquid phase separation (LLPS) of proteins (Burke et al., 2015; Lin et al., 2015; Molliex et al., 2015; Patel et al., 2015), while high concentrations can inhibit LLPS (Banerjee et al., 2017; Maharana et al., 2018). Conversely, many RNAs are capable of protein-free self-assembly (Jain and Vale, 2017; Langdon et al., 2018; Neil et al., 2020; Van Treeck et al., 2018). *In vivo*, recent studies have shown that RNA may be acting to form a non-dynamic or structural phase within a variety of biomolecular condensates (Clemson et al., 2009; Neil et al., 2020; Trcek et al., 2020; Van Treeck et al., 2018).

In *Xenopus* oocytes, RNAs required for germ layer patterning of the embryo, including *vg1* mRNA, are transported to the vegetal cortex in large cytoplasmic ribonucleoprotein (RNP) granules (reviewed in Cabral and Mowry, 2020). Recent work from our laboratory has characterized these RNPs as biomolecular condensates, termed Localization bodies (L-bodies) (Neil et al., 2020). L-bodies are large, irregularly shaped biomolecular condensates which are very highly enriched for localized RNA. *In vivo*, L-bodies are comprised of a non-dynamic, RNA-containing phase enmeshed by a comparatively dynamic protein layer. However, the mechanisms underlying the formation and maintenance of the biophysical state of L-bodies are not known.

Proteomic analysis of L-bodies revealed strong enrichment for proteins containing RNA binding domains (RBDs), intrinsically disordered regions (IDRs), or both types of domains (Neil et al., 2020), a conserved feature of phase separated condensates (reviewed in Banani et al., 2017). Many roles have been described for proteins with IDRs in biomolecular condensates, but the function of ordered, multivalent interaction domains are less well understood. Given the striking non-dynamic state of L-body RNAs, we were particularly interested in characterizing the role of multivalent RBPs in L-body assembly and dynamics. For this, we focused on polypyrimidine tract binding protein 3 (PTBP3), a previously uncharacterized L-body protein that we show in this work to be highly enriched within L-bodies. PTBP3 is a paralog of the well-characterized RBP, PTBP1 (Yamamoto et al., 1999), and PTB binding motifs within *vg1* mRNA are known to be required for *vg1* mRNA localization (Cote et al., 1999; Lewis et al., 2004). In addition to their roles in RNA localization, PTB proteins, such as PTBP1 (hnRNPI), PTBP2 (nPTB), and PTBP3 (ROD1), are involved in many steps in RNA metabolism depending on their subcellular localization and binding partners, including splicing, polyadenylation, mRNA stability, and translation initiation (reviewed in Hu et al., 2018; Sawicka et al., 2008). *In vitro*, PTBP1 phase separates in the presence of its RNA ligand (Banani et al., 2016; Li et al., 2012). PTB proteins contain four RNA Recognition Motifs (RRMs), which each bind polypyrimidine-rich sequences in RNA (Oberstrass et al., 2005), making PTB an ideal tool to study multivalent interactions of well folded interaction domains.

In this work, we elucidate the role of RNA-RBP interactions in the recruitment and dynamics of components of biomolecular condensates using PTBP3, both *in vivo* and *in vitro*. First, we demonstrate that PTBP3 is a novel L-body RNA binding protein with moderate *in vivo* mobility and show that PTB-RNA binding is required for the enrichment and dynamics of both protein and RNA in L-bodies. Next, we show that recombinant PTBP3 phase separates *in vitro* into irregularly shaped, solid or gel-like condensates in an RNA-dependent manner. *In vitro*, as *in vivo*, RNA forms a stable phase and PTBP3 dynamics are driven by binding to the non-dynamic RNA. Finally, we show that while a single RNA-RRM interaction is sufficient to target PTBP3 to L-bodies *in vivo*, multivalent interactions between RNA and protein work in concert to tune the mobility of PTBP3 after enrichment in L-bodies. Taken together, our results indicate that sequence-specific RNA-RBP interactions regulate recruitment of both RNA and proteins into condensates, but that formation of the non-dynamic RNA phase is concentration-dependent, rather than sequence-dependent. Importantly, the strength and number of protein interactions with the RNA phase drive dynamics after enrichment, suggesting a role for multivalent RNA-RBP interactions in regulating the physical properties of biomolecular condensates.

## Results

### *Enrichment and dynamics of* LE *RNA in L-bodies requires PTB binding sites*

Enrichment of localized RNAs in L-bodies can be recapitulated by a minimal localization element (*LE* RNA) derived from sequences within the 3’ UTR of the *vg1* mRNA (Cote et al., 1999; Lewis et al., 2004; Neil et al., 2020). *LE* RNA contains two pairs of PTB binding sites and strongly enriches within L-bodies (Figs. 1A-C, S1A-B). To test the role of PTB binding in the enrichment of RNA in L-bodies, we employed a mutated form of *LE* RNA, termed *mut PTB LE* RNA. The *mut PTB LE* RNA contains three U to A point mutations in each of the four PTB binding sites, but it otherwise identical to the *LE* RNA (Fig. S1A), and has been shown to no longer bind to PTBP1 (Lewis et al., 2004). When microinjected into oocytes, *mut PTB LE* RNA is only slightly enriched in L-bodies and, unlike the *LE* RNA, is also observed throughout the oocyte cytoplasm (Fig. 1C-C’’). Likewise, as seen at higher magnification, while the *LE* RNA is highly enriched within L-bodies (Figs. 1D, S1C-C’’), enrichment of *mut PTB LE* RNA is significantly reduced (Fig. 1D’-D’’). The *mut PTB LE* RNA is significantly less enriched in L-bodies, but is still more enriched than a non-localizing control, *XBM* RNA, which is neither enriched nor excluded from L-bodies (Fig. 1E, Fig. S1D-E). This low, but significant, level of enrichment is likely due to binding of *mut PTB LE* RNA to other L-body RBPs, such as Vera, which bind to sites not affected by the *mut PTB LE* RNA mutations (Lewis et al., 2004). Taken together, these data demonstrate that PTB binding to the *LE* RNA is required for robust enrichment of the RNA into L-bodies.

**Figure 1:**
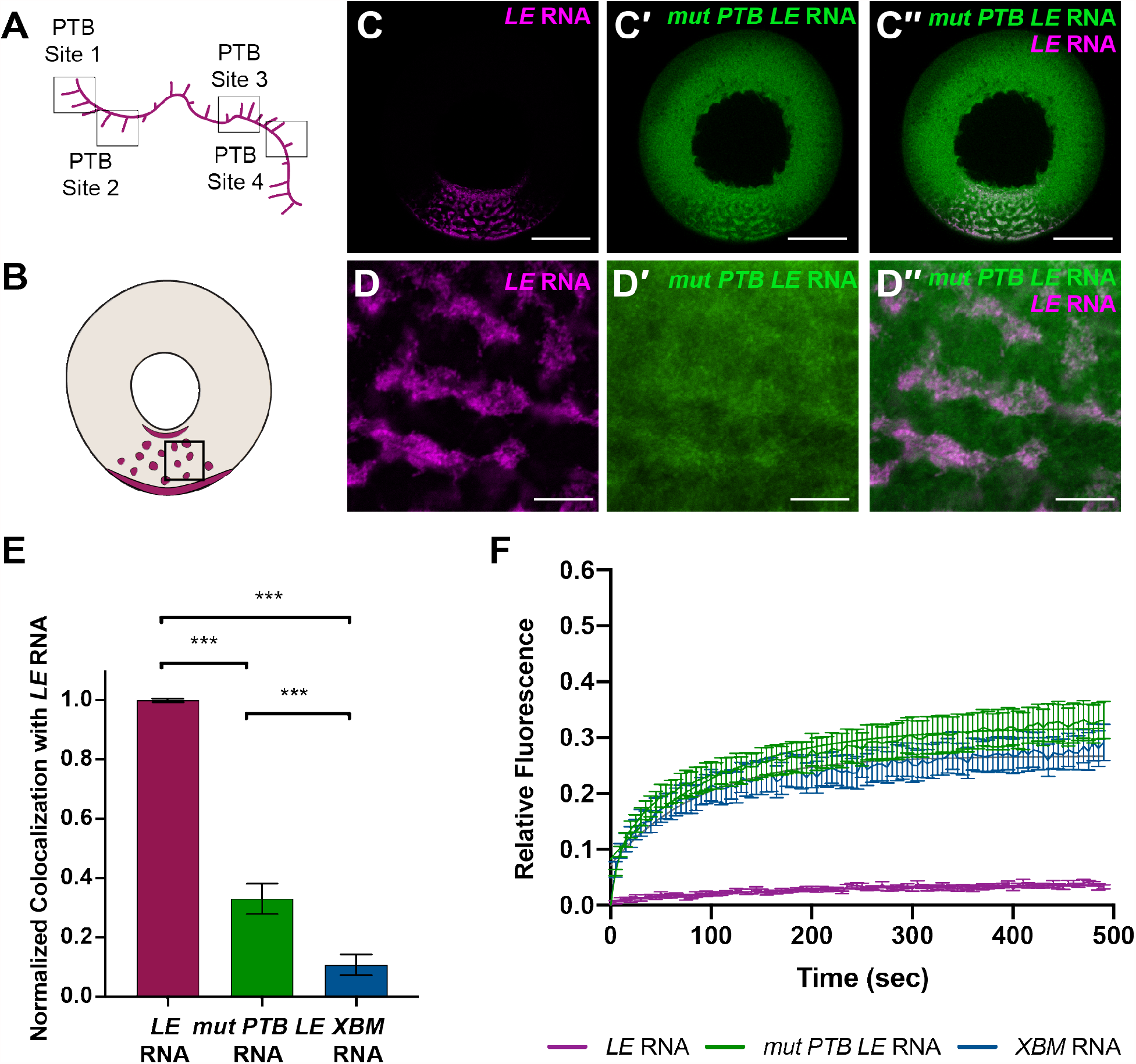
Enrichment and dynamics of *LE* RNA in L-bodies requires PTB binding sites. **(A)** Schematic of *LE* RNA (magenta) with four polypyrimidine tracts (PTB Sites 1-4) indicated in black boxes. **(B)** Schematic of a stage II *Xenopus* oocyte with LE RNA (magenta) localization as shown in whole oocyte images (as in C); the vegetal cortex is at the bottom. The portion of cytoplasm shown in high magnification images (as in D) is denoted by a black box. **(C)** Stage II oocytes were microinjected with fluorescently-labeled *LE* RNA (C, magenta) and *mut PTB LE* RNA, with all PTB binding sites mutated, (C’, green). The overlap is shown in C’’; scale bars=100 µm. **(D)** High magnification view of L-bodies in a stage II oocyte microinjected with *LE* RNA (D, magenta) and *mut PTB LE* RNA (D’, green). The overlap is shown in D’’; scale bars=10 µm. **(E)** Normalized Pearson correlation coefficient of Cy5-labeled *LE* RNA (magenta), *mut PTB LE* RNA (green), and *XBM* RNA (blue) with Cy3-labeled *LE* RNA in stage II oocytes, as in Fig. S1A-A’’, Fig. 1C-C’’, and Fig. S1C-C’’. *LE* RNA colocalization with *LE* RNA is set to 1. n=30 oocytes per RNA and error bars represent standard error of the mean. *** indicates p<0.01 **(F)** Stage II oocytes were microinjected with Cy5-labelled *LE* RNA to mark L-bodies, along with either Cy3-labelled *LE* RNA (magenta), *mut PTB LE* RNA (green) or *XBM* RNA (blue). Normalized FRAP recovery curves are shown. n=21 oocytes and error bars represent standard error of the mean.

As *LE* RNA is known to be non-dynamic in L-bodies (Neil et al., 2020), we next tested the dynamics of the *mut PTB LE* RNA in L-bodies. We microinjected oocytes with Cy3-labelled *LE* RNA, *mut PTB LE* RNA, or *XBM* RNA, along with Cy5-labelled *LE* RNA to mark L-bodies, and performed fluorescence recovery after photobleaching (FRAP) on the Cy3-labelled RNAs in L-bodies (Fig. 1F). As expected, *LE* RNA is almost entirely non-dynamic within L-bodies, with an immobile fraction of 93.9% (Fig. S1F). By contrast, *mut PTB LE* RNA is significantly more dynamic than the *LE* RNA, with an immobile fraction of 67.7%, and was indistinguishable from *XBM* RNA (72.4% immobile fraction). These results indicate that PTB binding sites are required for both enrichment of *LE* RNA in L-bodies and for the nondynamic nature of the RNA in L-bodies.

### PTBP3 enrichment and dynamics in L-bodies requires RNA binding

As PTB binding sites in *LE* RNA are necessary for RNA enrichment, we next tested if RNA binding by PTB is required for protein enrichment in L-bodies. L-bodies contain two paralogs of PTB, PTBP1 and PTBP3 (Neil et al., 2020). First, we tested the distribution of PTBP1 in oocytes by expressing mCherry tagged PTBP1 (mCh-PTBP1) in oocytes and assaying the subcellular localization via anti-mCh immunofluorescence (IF) (Fig. S2). We found that PTBP1, which shuttles between the nucleus and cytoplasm of *Xenopus* oocytes (Lewis et al., 2008; Xie et al., 2003), is only slightly enriched within L-bodies in stage II-III oocytes. Interestingly, unlike its paralog PTBP1, mCh-PTBP3 was not detected in the nucleus and is highly enriched in L-bodies (Fig. 2B). Therefore, we focused on PTBP3 for our experiments.

**Figure 2:**
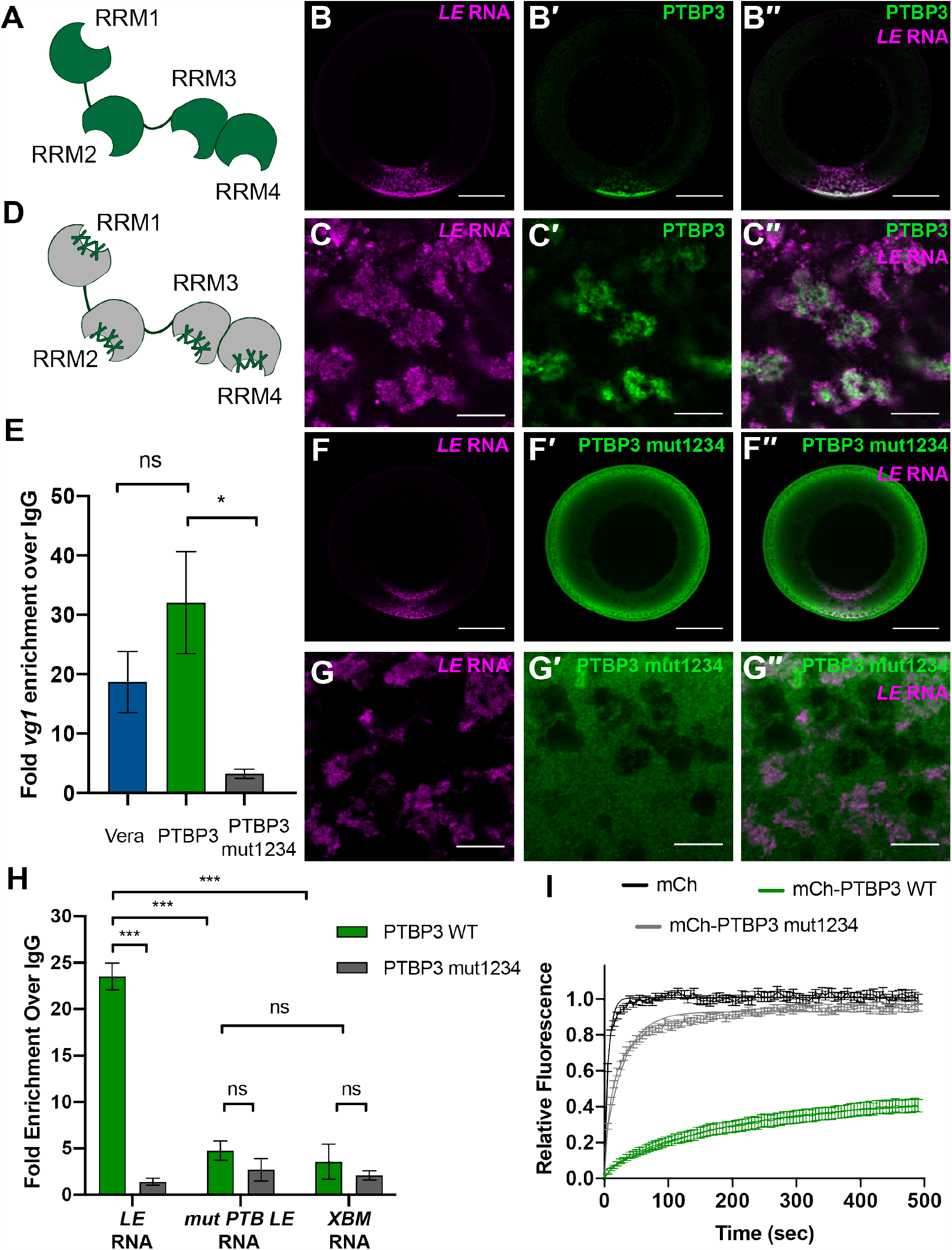
PTBP3 enrichment and dynamics in L-bodies require RNA binding. **(A)** Schematic of PTBP3 (green) with four RRMs (RRM1-4). **(B)** Fluorescently-labeled *LE* RNA (B, magenta) was microinjected into stage II oocytes expressing mCh-PTBP3, as detected by anti-mCh IF (B’, green). The overlap is shown in B’’; scale bar=100 µm. **(C)** High magnification view of L-bodies in a stage II oocyte microinjected with *LE* RNA (C, magenta) and expressing mCh-PTBP3, as detected by anti-mCh IF (C’, green). The overlap is shown in C’’; scale bar=10 µm. **(D)** Schematic of PTBP3 mut1234 (grey) with three point mutations (green X marks) introduced into each of the four RRMs (Table S1). **(E)** Lysates prepared from oocytes expressing Vera-mCh, mCh-PTBP3, or mCh-PTBP3 mut1234 were immunoprecipitated using anti-mCh and IgG. Following isolation of bound RNAs, *vg1* mRNA was detected via qRT-PCR, with normalization to a *luciferase* extraction control. Shown is fold enrichment for *vg1* mRNA over the IgG control. n=3 and error bars represent standard error of the mean. ns indicates p> 0.5, * indicates p<0.5 **(F)** Fluorescently-labeled *LE* RNA (F, magenta) was microinjected into stage II oocytes expressing mCh-PTBP3 mut1234, as detected by anti-mCh IF (F’, green). The overlap is shown in F’’; scale bar=100 µm. **(G)** High magnification view of L-bodies in a stage II oocyte microinjected with *LE* RNA (G, magenta) and expressing mCh-PTBP3 mut1234, as detected by anti-mCh IF (G’, green). The overlap is shown in G’’; scale bar=10 µm. **(H)** Oocytes expressing mCh-PTBP3 (green) or mCh-PTBP3 mut1234 (grey) were microinjected with *LE, mut PTB LE*, and *XBM* RNAs. Oocyte lysates were immunoprecipitated using anti-mCh and IgG. Following isolation, bound RNAs were detected by qRT-PCR, with normalization to a *luciferase* RNA extraction control. Shown is fold enrichment for each anti-mCh IP over the IgG control. n=3 and error bars represent standard error of the mean. ns indicates p>0.5, *** indicates p<0.01 **(I)** Stage II oocytes expressing mCh (black), mCh-PTBP3 (green), or mCh-PTBP3 mut1234 (grey) were microinjected with Cy5-labelled *LE* RNA to mark L-bodies. Normalized FRAP recovery curves are shown. n=21 oocytes and error bars represent standard error of the mean.

PTBP3 is an RNA binding protein containing four well-folded RNA recognitions motifs (RRMs; Fig. 2A). The PTBP3 RRMs bind to polypyrimidine tracts in RNA, here termed PTB sites (Yamamoto et al., 1999). To visualize the distribution of PTBP3 in oocytes, we expressed mCherry-tagged PTBP3 (mCh-PTBP3) in stage II-III oocytes and assayed the subcellular localization using anti-mCh IF. At low magnification, mCh-PTBP3 is highly enriched in the vegetal cytoplasm of oocytes, particularly at the vegetal cortex and in the lower vegetal cytoplasm and is co-localized with *LE* RNA (Fig. 2B-B’’). At higher magnification, mCh-PTBP3 is observed to be highly enriched in L-bodies (Fig. 2C-C’’). As PTBP3 is highly enriched within L-bodies *in vivo*, PTB binding sites are required for *LE* RNA localization, and it has a clear, multivalent RBD domain structure, we selected PTBP3 as a case study to probe the role of RNA binding in the localization of proteins to L-bodies and their dynamics after enrichment.

To determine whether RNA-binding is required for localization of mCh-PTBP3 to L-bodies, point mutations analogous to those that have been demonstrated to disrupt RNA-binding in human PTBP1 (Kafasla et al., 2011) were introduced into *Xenopus* PTBP3 (Fig. 2D, Table S1). Three point mutations were engineered into each of the four mCh-PTBP3 RRMs to create a quadruple RRM mutant (mCh-PTBP3 mut1234). First, to test whether the RRM mutations affected interaction between PTBP3 and *vg1* RNA, we expressed mCh-PTBP3 or mCh-PTBP3 mut1234 in oocytes and performed RNA immunoprecipitation (RIP) experiments using Vera-mCh, a well-established *vg1* mRNA binding protein as a positive control (Lewis et al., 2004) (Fig. 2E). mCh-PTBP3 immunoprecipitated endogenous *vg1* mRNA comparably to Vera-mCh, showing that PTBP3 interacts with *vg1* mRNA in oocytes. Conversely, immunoprecipitation of *vg1* mRNA was significantly decreased in the mCh-PTBP3 mut1234 injected oocytes, demonstrating that the mutations inserted into PTBP3 significantly reduce binding to endogenous *vg1* mRNA *in vivo*.

We also used RIP experiments to determine whether mutation of the PTB binding sites in *mut PTB LE* RNA blocked binding to PTBP3. Oocytes were microinjected with either mCh-PTBP3 or mCh-PTBP3 mut1234, along with *LE* RNA, *mut PTB LE* RNA, and *XBM* RNA. Following immunoprecipitation with anti-mCh and IgG as a negative control, the fold enrichment of each RNA was assayed via RT-PCR (Fig. 2H). As with the endogenous *vg1* mRNA, mCh-PTBP3 strongly immunoprecipitated the *LE* RNA, while the mCh-PTBP3 mut1234 did not; further demonstrating that the PTBP3 RRM mutations disrupt RNA binding. Importantly, neither mCh-PTBP3 nor mCh-PTBP3 mut1234 immunoprecipitated the *mut PTB LE* or *XBM* RNAs significantly over IgG controls. We conclude from these results that PTBP3 binds to one or more of the PTB binding sites in the *LE* RNA.

To determine whether binding of PTBP3 to RNA is required for enrichment of PTBP3 in L-bodies, we tested the subcellular distribution of mCh-PTBP3 mut1234 in stage II-III oocytes. In contrast to wild-type mCh-PTBP3 (Fig. 2B-B’’), mCh-PTBP3 mut1234 is distributed throughout the oocyte cytoplasm, and is not co-localized with *LE* RNA in L-bodies (Fig. 2F-F’’). At high magnification, mCh-PTBP3 mut1234 is neither enriched nor excluded from L-bodies, but is nearly ubiquitous in the vegetal cytoplasm (Fig. 2G-G’’), in marked contrast to wild-type PTBP3 (Fig. 2C-C’’). These results indicate that the mutations inserted into PTBP3 mut1234 that block RNA binding also disrupt localization to L-bodies.

Our previous work suggested a model for L-body structure in which localizing RNAs form a non-dynamic phase enmeshed in a dynamic protein phase in which the L-body proteins exhibit a range of moderate to high mobilities (Neil et al., 2020). Here, we hypothesize that the dynamics of L-body proteins are regulated by direct binding to the RNA phase. Thus, PTBP3, which we have shown binds *vg1* mRNA (Fig. 2E), should be moderately dynamic within L-bodies, while the quadruple RRM mutant should be significantly more dynamic. To test this hypothesis, we expressed mCh, mCh-PTBP3, or mCh-PTBP3 mut1234 in stage II oocytes and analyzed the protein dynamics *in vivo* using FRAP (Fig. 2I). As predicted, mCh-PTBP3 is moderately dynamic, with an immobile fraction of 52.4%, while mCh-PTBP3 mut1234 was significantly more dynamic, with an immobile fraction of 6.3%, and more closely resembled the mCh alone control. Taken together, these data indicate that binding of PTBP3 to RNA is necessary for its enrichment within L-bodies, and that RNA binding regulates the dynamics of the protein within L-bodies.

### *Recombinant PTBP3 and* LE *RNA phase separate into solid or gel-like condensates* in vitro

To probe the potential role of interactions between PTBP3 and RNA in phase separation, we developed an *in vitro* assay using recombinant PTBP3 (Fig. S3A) and *in vitro* transcribed *LE* RNA. Purified PTBP3 was mixed with *LE* RNA and incubated at room temperature for 1 hour without a crowding agent. To quantify the degree of phase separation across different conditions, turbidity was assayed as a measure of *in vitro* phase separation (Sanulli and Narlikar, 2021). First, to assess the role of electrostatic interactions on phase separation, the turbidity of PTBP3 and *LE* RNA or PTBP3 alone was tested across increasing salt concentrations (Fig. 3A). Across all NaCl concentrations tested, no significant turbidity was detected with PTBP3 alone, indicating that RNA is required for phase separation *in vitro*. However, with PTBP3 and *LE* RNA, turbidity was detected in a salt dependent manner; maximum phase separation was observed at intermediate levels of NaCl, but was inhibited at both low and high concentrations. Next, to assess the role of RNA on phase separation, the turbidity of PTBP3 and *LE* RNA or *LE* RNA alone was tested across an RNA concentration series (Fig. 3B). No phase separation was observed with *LE* RNA alone even at the highest concentrations tested, indicating that the observed phase separation requires PTBP3 and is not due solely to RNA-RNA interactions. With both PTBP3 and *LE* RNA, phase separation was detected in an RNA-dependent manner, indicating that RNA is required for phase separation. Importantly, phase separation is inhibited by high concentrations of *LE* RNA, indicating that multivalent interactions between PTBP3 and *LE* RNA may be important for *in vitro* condensate formation.

**Figure 3:**
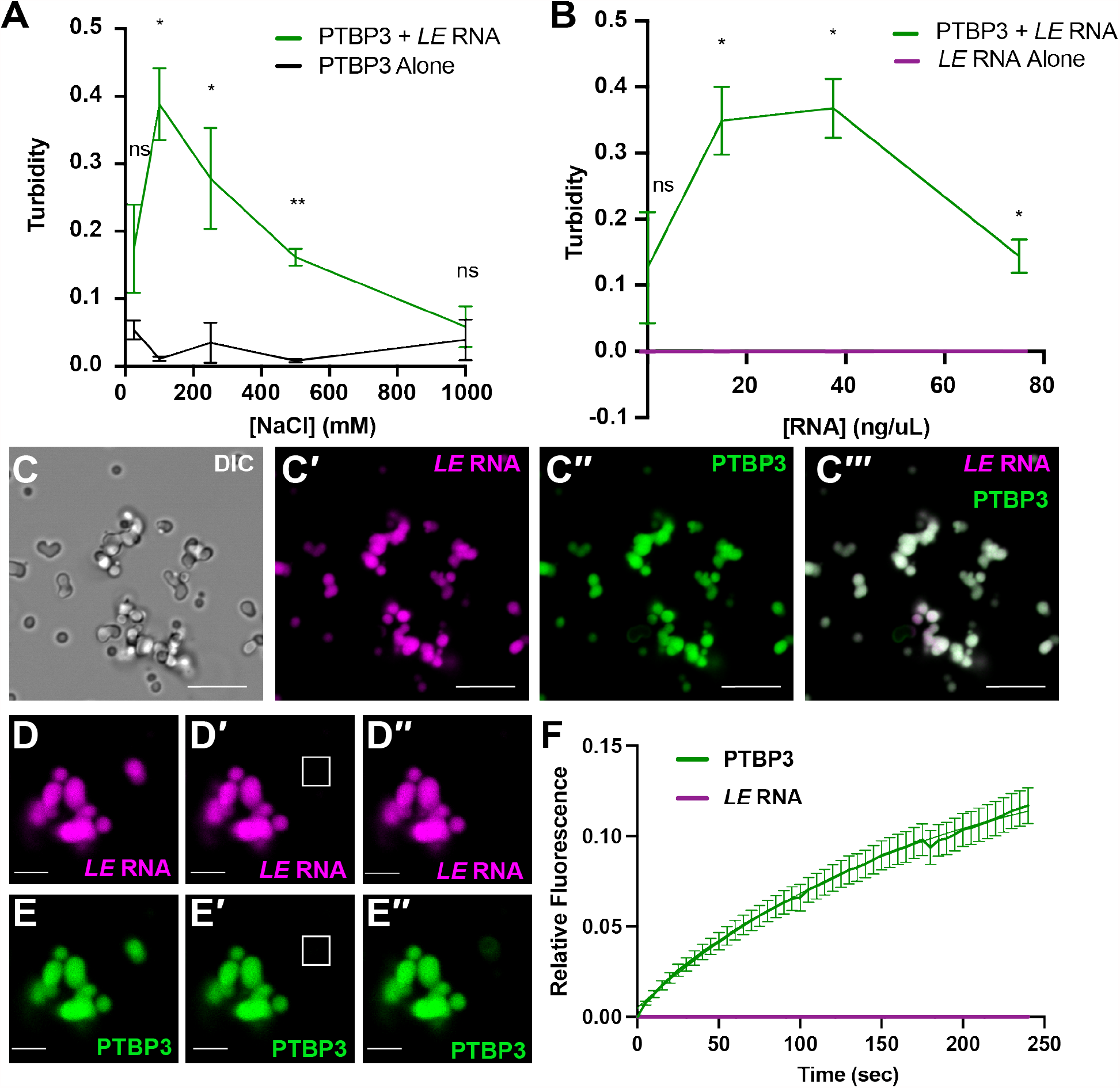
Recombinant PTBP3 and *LE* RNA phase separate into solid or gel-like condensates *in vitro*. **(A)** PTBP3 and *LE* RNA together (green) or PTBP3 alone (black) were incubated in the indicated concentrations of NaCl for 1 hr. at room temperature. Phase separation was monitored by turbidity, measured by OD600. Error bars represent the standard error of the mean, *n*=3. ns indicates p>0.5, * indicates p<0.5, ** indicates p<0.1 **(B)** PTBP3 and *LE* RNA together (green) or *LE* RNA alone (magenta) were incubated in the indicated concentrations of *LE* RNA for 1 hr. at room temperature. Phase separation was monitored by turbidity, measured by OD600. Error bars represent the standard error of the mean, *n*=3. ns indicates p>0.5, * indicates p<0.5. **(C)** AF647-labelled PTBP3 was incubated with AF488-labelled *LE* RNA for 1 hr. at room temperature. The resulting phase separated condensates are shown by DIC (C), and fluorescent imaging for *LE* RNA (C’, magenta) and PTBP3 (C’’, green). The overlay is shown in C’’’; scale bars=10 µm. **(D-E)** An image of PTBP3-*LE* condensates is shown with a 2 µm^2^ FRAP ROI (white). (D) *LE* RNA fluorescence pre-bleach, post-bleach (D’), and 4 min. post-recovery (D’’). Scale bars=2 µm. (E) AF647-PTBP3 fluorescence pre-bleach, post-bleach (E’), and 4 min. post-recovery (E’’). **(F)** Normalized FRAP recovery curves are shown for PTBP3-*LE* RNA condensates, carried out as in D-E. *n*=21 condensates and error bars represent standard error of the mean.

To further analyze *in vitro* phase separation of PTBP3 and *LE* RNA, the condensates formed at intermediate salt and RNA concentrations were visualized by microscopy. Under these conditions, PTBP3 and *LE* RNA formed irregularly shaped structures that were highly enriched for both the protein and RNA components (Fig. 3C-C’’’). While some condensates appeared round, as would be expected for liquid condensates due to surface tension (Hyman et al., 2014), other condensates appeared to be comprised of multiple small droplets that had interacted to form larger structures, but not fused and reformed into a larger sphere. To characterize the dynamics of the protein and RNA in the *in vitro* phase separated condensates, we performed FRAP. *LE* RNA (Fig. 3D-D’’) was non-dynamic *in vitro* (immobile fraction=99.8%), indicating that in *in vitro* phase separated condensates, *LE* RNA forms a solid or gel-like phase (Fig. 3F, S3B). PTBP3 (Fig. 3E-E’’), while significantly more dynamic than *LE* RNA, was only moderately dynamic within the *in vitro* phase separated condensates (immobile fraction=82.0%) (Fig. 3F, S3B). These results demonstrate that interactions between *LE* RNA and PTBP3 are sufficient to drive phase separation into irregularly shaped condensates *in vitro*, with trends in RNA and protein dynamics mirroring the findings in *in vivo* L-bodies: both *in vivo* and *in vitro, LE* RNA forms a non-dynamic phase and PTBP3 is moderately dynamic.

### In vitro *condensate morphology and PTBP3 dynamics are dependent on RNA-binding*

To test whether phase separation of PTBP3 *in vitro* is dependent on specific RNA sequences, we compared the phase separation of PTBP3 in the presence of *mut PTB LE* RNA, *XBM* RNA, and *LE* RNA, as above. For each of the RNAs, PTBP3 formed phase separated condensates that incorporated both PTBP3 and the RNA (Fig. 4A-C), demonstrating that although *in vitro* phase separation requires RNA, it does not require interaction of PTBP3 specifically with PTB sites, which are present only in *LE* RNA. For *in vitro* condensates, unlike *in vivo* L-bodies, there was no difference in partitioning of *LE* RNA, *mut PTB LE* RNA, or *XBM* RNA into PTBP3 condensates (Fig. S4A). However, the morphologies of the condensates formed by PTBP3 with *LE* RNA, *mut PTB LE* RNA, and *XBM* RNA are distinct from one another (Fig. S4C-E). PTBP3 and *LE* RNA (Fig. 4A, S4C-E) formed fewer, larger, and more irregularly shaped condensates, whereas condensates formed with PTBP3 and *XBM* RNA were smaller, more numerous, and significantly more circular (Fig. 4C, S4C-E). Condensates formed with PTBP3 and *mut PTB LE* RNA displayed an intermediate morphology, but more closely resembled *XBM* RNA condensates (Fig. 4B, S4C-E). Importantly, none of these differences were due to differences in RNA size as all *in vitro* transcribed RNAs were precisely length matched.

**Figure 4:**
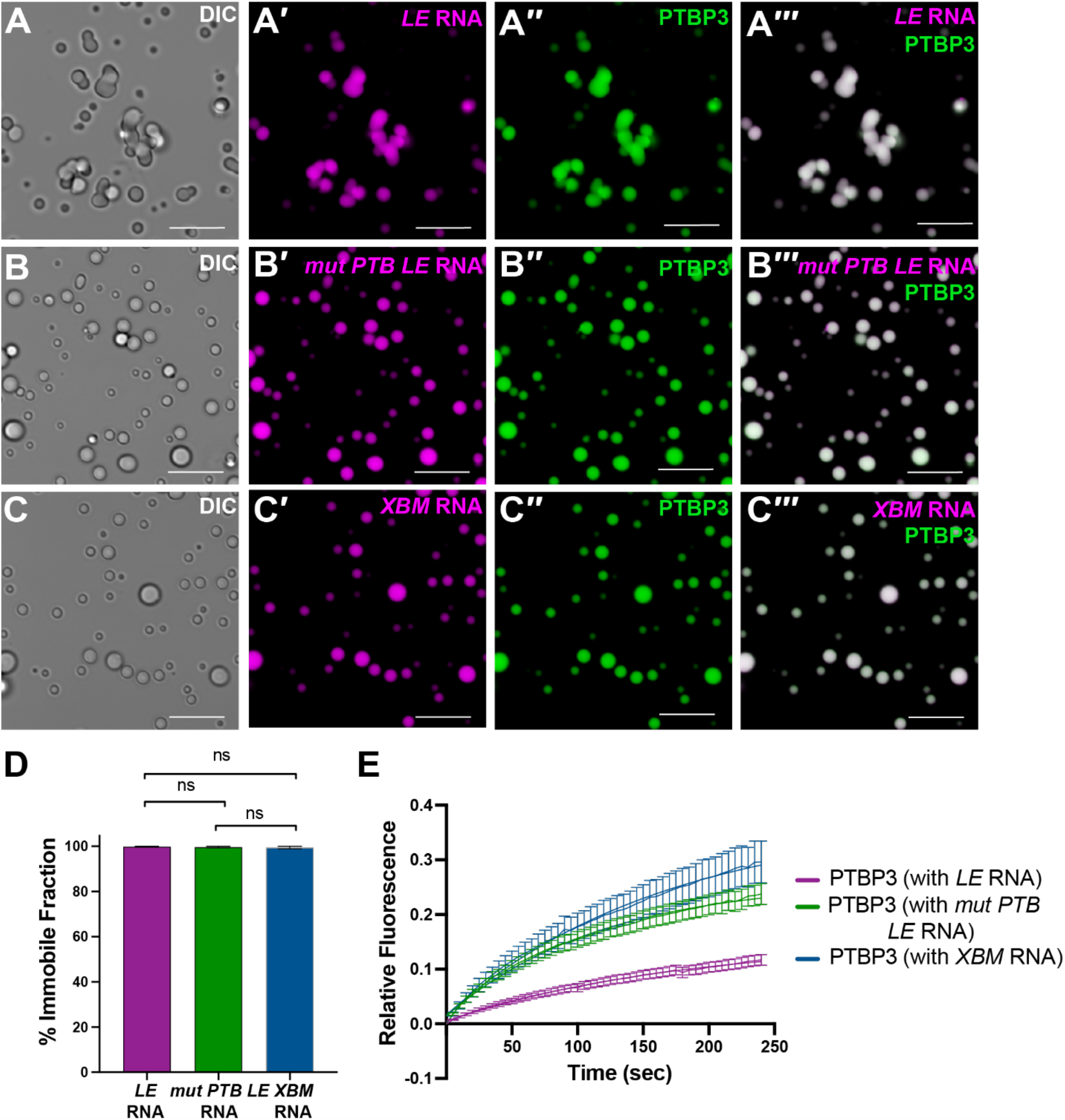
*In vitro* condensate morphology and PTBP3 dynamics are dependent on RNA-binding. **(A)** AF647-labelled PTBP3 was incubated with AF488-labelled *LE* RNA for 1 hour at room temperature. The resulting phase separated condensates are shown in DIC (A), and by fluorescence imaging for *LE* RNA (A’, magenta), and PTBP3 (A’’, green). The overlay is shown in A’’’; scale bars=10 µm. **(B)** AF647-labelled PTBP3 was incubated with AF488-labelled *mut PTB LE* RNA for 1 hr. at room temperature. The resulting phase separated condensates are shown in DIC (B), and by fluorescence imaging for *mut PTB LE* RNA (B’, magenta), and PTBP3 (B’’, green). The overlay is shown in B’’’; scale bars=10 µm. **(C)** AF647-labelled PTBP3 was incubated with AF488-labelled *XBM* RNA for 1 hr. at room temperature. The resulting phase separated condensates are shown in DIC (C), and by fluorescence imaging for *XBM* RNA (C’, magenta), and PTBP3 (C’’, green). The overlay is shown in C’’’; scale bar=10 µm. **(D)** Percent immobile fraction for RNA FRAP of *in vitro* condensates. Error bars represent standard error of the mean. ns indicates p>0.5 **(E)** Normalized FRAP recovery curves for are shown for PTBP3 in condensates containing PTBP3-*LE* RNA (magenta), PTBP3-*mut PTB LE* RNA (green), and PTBP3-*XBM* RNA (blue). *n*=21 condensates per RNA and error bars represent standard error of the mean.

The differences in condensate morphology led us to hypothesize that these different types of condensates may have varying protein and/or RNA dynamics. To test this, we performed FRAP on the protein and RNA in each type of *in vitro* condensate. Surprisingly, each of the RNAs was non-dynamic, exhibiting mobilities that are not significantly different from one another (Fig. 4D), with immobile fractions for whole condensate FRAP of 99.8%, 99.7%, and 99.5% for the *LE, mut PTB LE*, and *XBM* RNA, respectively. Similarly, all RNAs were also non-dynamic via partial condensate FRAP (Fig. S5). These data suggest that *in vitro*, as in L-bodies *in vivo*, RNA forms a non-dynamic phase. However, unlike *in vivo*, this non-dynamic RNA phase forms *in vitro* regardless of the primary sequence of the RNA. Conversely, dynamics of PTBP3 protein were dependent on the sequence of the RNA incorporated into the condensates; PTBP3 was least mobile in condensates formed with *LE* RNA (immobile fraction=82.0%), and significantly more mobile in condensates formed with *mut PTB LE* (immobile fraction=71.5%) or *XBM* RNA (immobile fraction=64.8%) (Fig. 4E, Fig. S4B). These data suggest that PTBP3 dynamics *in vitro* are dependent on the ability to interact with the non-dynamic RNA, consistent with our *in vivo* results.

### PTBP3 dynamics in L-bodies are dependent on multivalent interactions with RNA

Because PTBP3 dynamics depend on interaction with RNA both *in vivo* and *in vitro*, we next asked whether this relies on specific PTBP3 RRMs or combinations of PTBP3 RRMs. To test the role of multivalency, we took an *in vivo* approach as recombinant PTBP3 RRM mutants exhibited poor solubility *in vitro*, particularly after cleavage of the MBP tag. However, *in vivo*, the expression and solubility of the mutant proteins in oocytes were comparable to that of WT PTBP3 (Fig. S6). First, we engineered PTBP3 double RRM mutants (mut12 and mut34), expressed them in stage II oocytes, and tested the double RRM mutants for their ability to bind *LE* RNA *in vivo* by RIP (Fig. 5A). mCh-PTBP3 mut12 immunoprecipitated *LE* RNA comparably to the wild type protein, suggesting that RRMs 1 and 2 are not required for binding *LE* RNA *in vivo*. By contrast, mCh-PTBP3 mut34 immunoprecipitated *LE* RNA comparably to mCh-PTBP3 mut1234, suggesting that RRM3, RRM4, or both RRMs are required for binding to *LE* RNA. To distinguish between these possibilities, we created single RRM mutants (mut3 and mut4) and tested their ability to immunoprecipitate *LE* RNA. Both mCh-PTBP3 mut3 and mCh-PTBP3 mut4 immunoprecipitated *LE* RNA comparably to the wild type and mCh-PTBP3 mut12 proteins, demonstrating that *LE* RNA can bind to RRM3 and RRM4. Hence, intact forms of RRM3 or RRM4 are both necessary and sufficient for robust *LE* RNA binding *in vivo*.

**Figure 5:**
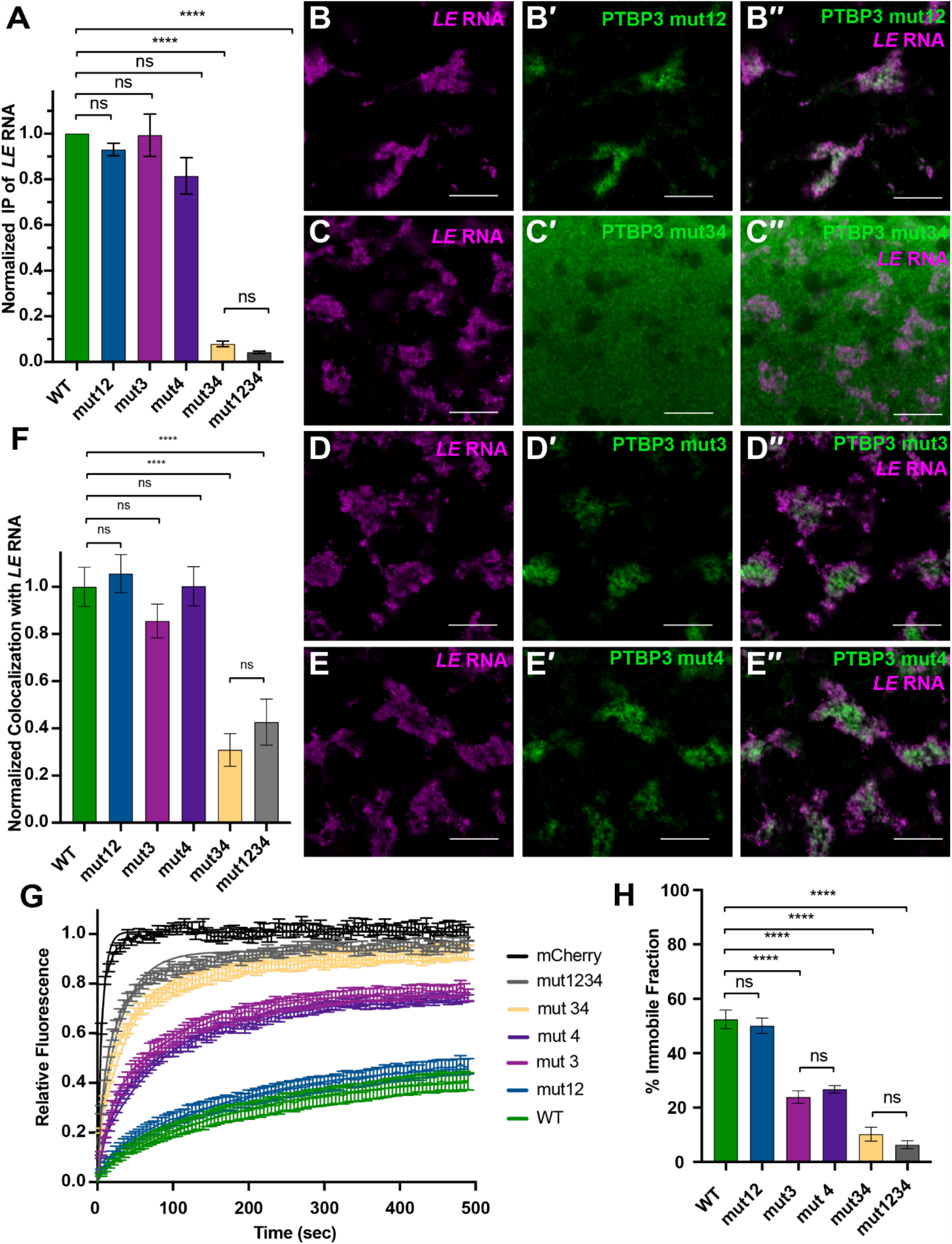
PTBP3 dynamics in L-bodies are dependent on multivalent interactions with RNA. **(A)** Oocytes expressing mCh-PTBP3 (green), mCh-PTBP3 mut12 (blue), mCh-PTBP3 mut34 (yellow), mCh-PTBP3 mut3 (magenta), mCh-PTBP3 mut4 (purple), and mCh-PTBP3 mut1234 (grey) were microinjected with *LE* RNA. Oocyte lysates were immunoprecipitated using anti-mCh and IgG. Following isolation of bound RNAs, *LE* RNA was detected via qRT-PCR, with normalization to a luciferase RNA extraction control. Fold enrichment for mCh-PTBP3 WT over the IgG control is set to 1. n=3 and error bars represent standard error of the mean. ns indicates p>0.5, **** indicates p<0.001. **(B)** High magnification view of L-bodies in a stage II oocyte microinjected with LE RNA (B, magenta) and expressing mCh-PTBP3 mut12, as detected by anti-mCh IF (B’, green). The overlap is shown in B’’; scale bars=10 µm. **(C)** High magnification view of L-bodies in a stage II oocyte microinjected with *LE* RNA (C, magenta) and expressing mCh-PTBP3 mut34, as detected by anti-mCh IF (C’, green). The overlap is shown in C’’; scale bars=10 µm. **(D)** High magnification view of L-bodies in a stage II oocyte microinjected with *LE* RNA (D, magenta) and expressing mCh-PTBP3 mut3, as detected by anti-mCh IF (D’, green). The overlap is shown in D’’; scale bars=10 µm. **(E)** High magnification view of L-bodies in a stage II oocyte microinjected with *LE* RNA (E, magenta) and expressing mCh-PTBP3 mut4, as detected by anti-mCh IF (E’, green). The overlap is shown in E’’; scale bars=10 µm. **(F)** Normalized Pearson correlation coefficient of mCh-PTBP3 WT (green), mCh-PTBP3 mut12 (blue), mCh-PTBP3 mut34 (yellow), mCh-PTBP3 mut3 (magenta), mCh-PTBP3 mut4 (purple), and mCh-PTBP3 mut1234 (grey) with *LE* RNA in stage II oocytes. mCh-PTBP3 WT colocalization with *LE* RNA is set to 1. n=30 oocytes per protein and error bars represent standard error of the mean. ns indicates p>0.5, **** indicates p<0.001 **(G)** Stage II oocytes expressing mCh-PTBP3 WT (green), mCh-PTBP3 mut12 (blue), mCh-PTBP3 mut34 (yellow), mCh-PTBP3 mut3 (magenta), mCh-PTBP3 mut4 (purple), and mCh-PTBP3 mut1234 (grey) were microinjected with Cy5-labelled *LE* RNA to mark L-bodies. Normalized FRAP recovery curves are shown. n=21 oocytes and error bars represent standard error of the mean. **(H)** Percent immobile fraction for protein FRAP experiments shown in (G). Error bars represent standard error of the mean. ns indicates p>0.5, **** indicates p<0.001.

To test whether specific PTBP3 RRMs drive enrichment into L-bodies, we expressed each of the double and single PTBP3 RRM mutants *in vivo* and assayed their subcellular distribution using mCh IF. We found that *LE* RNA binding by PTBP3 RRM mutants predicts enrichment within L-bodies: mCh-PTBP3 mut12 is strongly enriched within L-bodies (Fig. 5B-B’’,F), while mCh-PTBP3 mut34 showed no enrichment (Fig. 5C-C’’,F); mCh-PTBP3 mut3 and mCh-PTBP3 mut4 single RRM mutants strongly enriched within L-bodies (Fig. 5D-D’’, E-E’’,F). These results show that binding to RNA by either RRM3, RRM4, or RRMs 3 and 4 is sufficient to drive localization and enrichment of PTBP3 in L-bodies and that this enrichment does not require multivalent interactions between protein and RNA.

While RNA binding by a single RRM is sufficient to drive protein localization, we reasoned that interaction between the non-dynamic RNA phase and multiple RRMs may be required for the moderate dynamics of the wild type PTBP3 protein. To test the role of multivalent interactions in protein dynamics, we tested each of the PTBP3 mutants by FRAP (Fig. 5G-H). The dynamics of mCh-PTBP mut12 (immobile fraction=50.1%), was indistinguishable from the wild type protein, and mCh-PTBP3 mut34 (immobile fraction=10.2%) was indistinguishable from mCh-PTBP3 mut1234, as in the RIP and colocalization experiments. However, as predicted by our model, mCh-PTBP3 mut3 and mCh-PTBP3 mut 4 showed intermediate dynamics with immobile fractions of 23.9% and 26.7%, respectively, and were indistinguishable from one another. These results demonstrate that while RNA-PTBP3 binding to a single RRM is sufficient for enrichment in L-bodies, multivalent interactions with RNA act in combination to regulate the dynamics of the protein within the L-body.

## Discussion

### Proposed model for L-body component recruitment and dynamics

In this work, we have dissected the role of RBP-RNA binding in L-bodies using both *in vivo* and *in vitro* techniques. L-bodies are newly identified, irregularly shaped biomolecular condensates with a non-dynamic RNA phase and a comparatively dynamic protein phase (Neil et al., 2020). However, the mechanisms underlying the range of biophysical states observed in L-body components were unclear. Here, we propose a multistep model for L-body component recruitment and dynamics based on both specific RNA-RBP binding and RNA concentration-dependent effects (Fig. 6).

**Figure 6:**
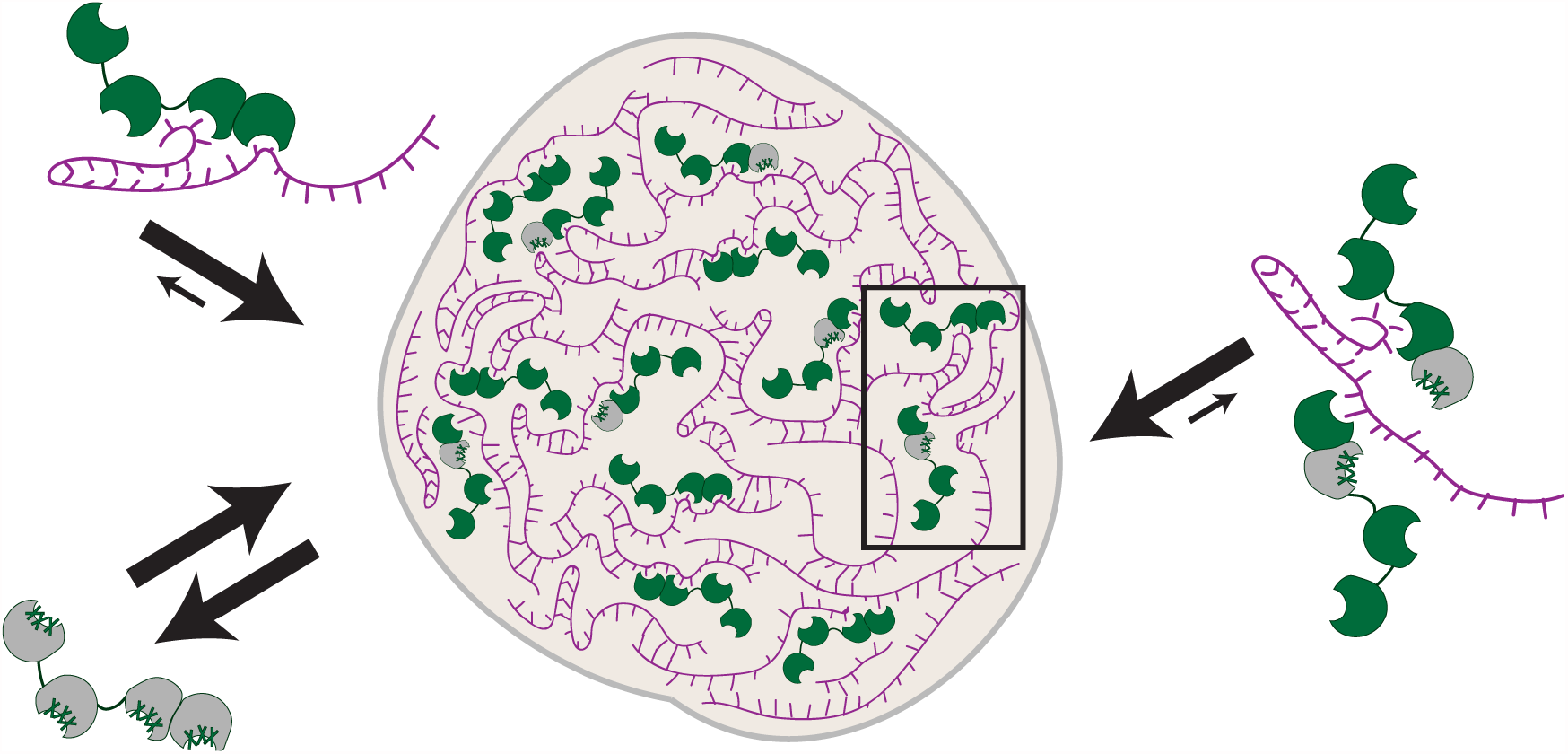
Model for L-body recruitment and dynamics. RBPs, such as PTBP3 (green), bind localizing RNAs (magenta) in a sequence-dependent manner which is required for both RNA and protein enrichment in L-bodies (tan), such that PTBP3 RNA-binding mutants (grey) are neither enriched nor excluded from L-bodies. Once enriched, localizing RNAs form a solid or gel-like phase in a sequence-independent manner within the L-body. RNA-protein interaction with one RRM is sufficient to target PTBP3 to L-bodies, while the strength of number of interactions with non-dynamic RNA tunes the protein dynamics within the L-body (black rectangle).

First, RBPs bind localizing RNAs in the oocyte cytoplasm, driving the enrichment of the RBP and RNA within L-bodies. Accordingly, PTBP3 strongly enriches in L-bodies, but PTBP3 mut1234, which no longer binds to the *LE* RNA, is nearly ubiquitous throughout the cytoplasm. Similarly, *LE* RNA enrichment is a defining feature of L-bodies, but *mut PTB LE* and *XBM* RNAs, which do not bind to PTB, are both distributed throughout the cytoplasm. Therefore, sequence specific RNA-protein binding is required for both the RNA and the RBP to enrich within L-bodies in *Xenopus* oocytes. The *in vitro* RNA-PTBP3 condensates, however, enrich equally for all RNAs as this step of regulation is not present in the minimal *in vitro* system. Instead, non-specific interactions between PTBP3 and *mut PTB LE* or *XBM* RNA are sufficient to drive phase separation *in vitro*.

Next, following RBP and RNA enrichment, locally high concentrations of RNA within the L-body facilitate intermolecular RNA-RNA interactions, which leads to the formation of a solid or gel-like RNA phase. *In vivo, LE* RNA, which is highly concentrated within the L-body, is almost entirely nondynamic, while *mut PTB LE* and *XBM* RNAs are not enriched within L-bodies and are much more dynamic. These dynamics are not a property of RNA size, as *LE, mut PTB LE*, and *XBM* RNAs are precisely length-matched and previous work has demonstrated that RNA mobility in *in vivo* L-bodies also does not correlate with RNA length (Neil et al., 2020). While the *in vivo* data do not distinguish between differences in RNA dynamics due to lack of PTBP3 binding and RNA concentration dependent effects, the *in vitro* data suggests that the process is RNA concentration dependent; RNA gelation occurs in the *in vitro* PTBP3-RNA condensates, driving *LE, mut PTB LE*, and *XBM* RNAs to all form non-dynamic phases regardless of PTB binding.

Finally, it is both the strength and number of interactions with the RNA phase that tune protein dynamics. In oocytes, a single PTBP3 RRM-RNA interaction is sufficient to drive enrichment of PTBP3 into L-bodies, but multivalent interactions between multiple RRMs and the RNA work in combination to regulate the mobility of the protein after enrichment. In accordance with this idea, another direct *LE* RNA binding protein, Vera, was also found to be only moderately dynamic *in vivo*, with an immobile fraction of 47.1% (Neil et al., 2020). *In vitro*, it is the strength of the interaction between PTBP3 and the RNA phase that determines both condensate morphology and PTBP3 dynamics: PTBP3-*LE* RNA condensates are irregularly shaped and have the lowest PTBP3 dynamics, while PTBP3-*XBM* RNA condensates are more circular and have the highest PTBP3 dynamics.

### Non-dynamic RNA phases facilitate the formation of irregularly shaped condensates

In addition to L-bodies, recent studies have identified other non-spherical biomolecular condensates, including TIS granules which form through the phase separation of an RBP, TIS11B, near the endoplasmic reticulum (ER) (reviewed in Fare et al., 2021; Ma and Mayr, 2018). The existence of irregularly shaped condensates is somewhat counterintuitive as the effects of surface tension drive many liquid-like condensates to be spherical in shape (Hyman et al., 2014). However, non-dynamic RNAs may be a conserved mechanism to build condensates with irregular morphologies that still have some liquid-like properties, such as highly dynamic proteins.

*In vitro* studies have provided insights into the mechanisms underlying the irregular condensate morphologies observed. Here, we have shown the PTBP3 and *LE* RNA phase separate *in vitro* into irregularly shaped condensates. However, PTBP3 also phase separates into more spherical condensates in the presence of RNAs without clear PTB binding sites, including the *mut PTB LE* and *XBM* RNAs, indicating that specific RNA-RBP interactions may drive the formation of a more gel-like condensate. In all cases, RNA forms a non-dynamic phase *in vitro* which we hypothesize to be due to concentration-dependent rather than sequence-specific effects. Recent *in vitro* studies using a fusion protein of the RBD of TIS11B and the IDR of FUS (FUS-TIS) and various RNAs also produced condensates with varying morphologies (Ma et al., 2021). However, rather than observing morphological differences based on specific RNA-RBD binding as seen with PTBP3, FUS-TIS condensate morphology varied based on the ability of the RNA to form intermolecular RNA-RNA interactions. Additionally, RNAs in spherical FUS-TIS condensates were found to be dynamic via FRAP, while RNAs in irregularly shaped condensates were nondynamic. The differences between the *in vitro* condensates may be due to variation in the rate of aging of the condensates *in vitro* or differences in the type of protein tested-PTBP3 is highly ordered and is comparatively non-dynamic via FRAP, while FUS-TIS contains a well-established IDR and is highly dynamic via FRAP. However, both studies demonstrate that a stable RNA phase formed by certain RNAs can lead to failed condensate fusion events *in vitro*, driving the formation of an irregularly shaped condensate.

Although the functions of irregularly shaped biomolecular condensates remain unclear *in vivo*, these morphologies increase the surface area to volume ratio, increasing the interaction interface with the cytoplasm. In L-bodies, this may be beneficial for efficiently capturing RNPs which have not yet been incorporated into L-bodies and are diffusing in the cytoplasm. As the later translation of RNAs incorporated into L-bodies is required to pattern the embryo, a high degree of enrichment of localizing RNAs may be required to prevent misexpression of these genes.

### Emergence of biomolecular condensates with non-dynamic RNA and dynamic proteins

Biomolecular condensates exist on a continuum of biophysical states from a demixed liquid to a solid state, often with different components of the condensate displaying varying dynamics (reviewed in Alberti et al., 2019; Fare et al., 2021). Recent studies have demonstrated a growing variety of biomolecular condensates which have non-dynamic RNAs and comparatively dynamic proteins, including *Drosophila* germ granules, paraspeckles, and *Xenopus* L-bodies (Mao et al., 2011; Neil et al., 2020; Trcek et al., 2020). As each of these types of biomolecular condensates is enriched for RNAs, intermolecular RNA-RNA interactions may be particularly thermodynamically favorable due to the high local concentration of RNA (Van Treeck and Parker, 2018), driving the formation of the non-dynamic RNA phase. The growing number of condensates with stable RNA phases suggests that intermolecular RNA-RNA interactions driving the formation of a stable RNA substructure may be a common feature of many biomolecular condensates, giving broad relevance to the insights into the role of RNA binding on protein dynamics in L-bodies.

In L-bodies, the strength and number of the interactions with the stable RNA phase determine PTBP3 protein dynamics. However, the role of other proteins, particularly proteins containing IDRs, in maintaining the biophysical state of the L-body is not yet understood. While *in vitro* PTBP3-RNA condensates and *in vivo* L-bodies contain non-dynamic RNA and moderately dynamic PTBP3, both *LE* RNA and PTBP3 were less dynamic *in vitro* than *in vivo*, suggesting that other factors contribute to the maintenance of the biophysical state of L-bodies *in vivo*. As L-bodies, like many other biomolecular condensates, are highly enriched for proteins with multiple RNA binding domains and IDRs, the relative contribution of each of these types of interactions on the dynamics of the condensate is an important outstanding question. For example, in FXR1 assemblies, RNA binding drives phase separation and IDRs of various lengths tune the dynamics of the condensate (Smith et al., 2020). Similarly, in L-bodies IDR-containing proteins may function to keep the condensate in a more liquid-like state, perhaps preventing an irreversible transition to a solid-like state. Additionally, L-bodies contain helicases and post-translational modifying enzymes (Neil et al., 2020), which may function to remodel interactions *in vivo*.

As additional biomolecular condensates continue to be identified across an ever-growing diversity of cell types and subcellular localizations, it remains an important challenge to characterize the principles which are conserved or divergent across many classes of condensates. One hallmark of biomolecular condensates is the enrichment for RNA and multivalent RNA binding proteins, highlighting the importance of understanding how RNAs, proteins, and RNA-protein interactions contribute to the characteristics of condensates. Our results, indicating that protein dynamics are tuned by multivalent interactions with a concentration-dependent RNA phase, are therefore likely to provide a general paradigm that is applicable to other classes of biomolecular condensates.

## Materials and Methods

### Oocyte Isolation and Culture

All animal work was approved by the Brown University Institutional Animal Care and Use Committee. Oocytes were surgically harvested from wild type *Xenopus laevis* females (Nasco, #LM00535MX), enzymatically defolliculated using 2 mg/mL collagenase (Sigma), and washed in MBSH [88 mM NaCl, 1 mM KCl, 2.4 mM NaHCO_3_, 0.82 mM MgSO_4_, 0.33 mM Ca(NO_3_)_2_, 0.41 mM CaCl_2_, 100mM HEPES (pH 7.6)]. Stage II-III oocytes were manually sorted, and cultured at 18°C in OCM+ [50% Leibovitz L-15 medium, 15 mM HEPES (pH 7.6), 1 mg/mL insulin, 50 U/mL nystatin (Sigma), 100 U/mL penicillin/streptomycin (ThermoFisher), 0.1 mg/mL gentamicin (ThermoFisher), 0.1 mg/mL ciprofloxacin] (Sigma).

### RNA transcription

For protein coding RNAs, RNAs were transcribed *in vitro* with the SP6 mMessage machine kit (Ambion) using the following linearized plasmids as the DNA template: pSP64:mCh-PTBP3 WT, pSP64:mCh-PTBP3 mut12, pSP64:mCh-PTBP3 mut34, pSP64:mCh-PTBP3 mut3, pSP64:mCh-PTBP3 mut4, pSP64:mCh-PTBP3 mut1234, pSP64:mCh, pSP64:mCh-PTBP1, and pSP64:Vera-mCh (Neil et al., 2020). RNAs were cleaned up by phenol-chloroform extraction and isopropanol precipitation as previously described (Jeschonek and Mowry, 2018). The concentration of the RNA was measured via Qubit RNA BR assay (ThermoFisher).

For the barcoded RNAs, RNAs were transcribed *in vitro* with the T7 MEGA script kit (Ambion) using length matched PCR products as the DNA templates. PCR products were generated from the following plasmids, with the forward primer for each construct containing the T7 promoter sequence and the unique barcode sequence as shown in Table S2: pSP73:2×135 (*LE)* (Gautreau et al., 1997), pSP73:2×135ΔVM1 (*mut PTB LE)* (Lewis et al., 2004), pSP64:XBM (*XBM)* (Krieg and Melton, 1984). To synthesize fluorescently labelled RNAs for oocyte microinjection, RNAs were transcribed as above in the presence of 250 nM Cy3 or Cy5-UTP (ThermoFisher). For fluorescently labelled RNAs for *in vitro* phase separation experiments, RNAs were transcribed as above in the presence of 50 nM AF488-UTP (ThermoFisher). T7 transcribed RNAs were cleaned up using MEGAclear transcription clean up kits (ThermoFisher) and the concentration of RNA was measure via Qubit RNA broad range assay (ThermoFisher).

### Whole mount immunofluorescence (IF) and colocalization analysis

Oocytes were microinjected with 2 nL of 500 nM RNA encoding an mCh-PTBP3 construct and 250 nM Cy-UTP labelled *LE* RNA to mark L-bodies. Oocytes were cultured for 48 hours in OCM+, fixed for 1 hour in fixation buffer [80 mM PIPES (pH 6.8), 1 mM MgCl_2_, 5 mM EGTA, 0.2% Triton X-100, 3.8% formaldehyde], and washed 3 times for 15 min. each in PBT [137 mM NaCl, 2.7 mM KCl, 10 mM Na_2_HPO_4_, 1.8 mM KH_2_PO_4_, 0.2% BSA, 0.1% Triton X-100]. Oocytes were blocked for 4 hours at room temperature in PBT+ [PBT supplemented with 2% goat serum and 2% BSA], incubated overnight at 4°C in a 1:500 dilution of anti-mCh primary antibody (Abcam, ab62341) in PBT+, and washed 3 times for 2 hours each in PBT. Oocytes were incubated overnight at 4°C in a 1:1000 dilution of goat, anti-rabbit AF647 conjugated secondary antibody (ThermoFisher, 21235), washed 3 times for 2 hours each, dehydrated in anhydrous methanol, and frozen at −20°C until imaging. Immediately prior to imaging, oocytes were cleared in BABB solution (1:2 benzyl alcohol:benzyl benzoate). Oocytes were imaged on an inverted Olympus FV3000 confocal microscope using 20× UPlan Super Apochromat objective (air, NA=0.75) and 60× UPlan Super Apochromat objective (silicon oil, NA=1.3) using GaAsP detectors.

For each of the mCh-PTBP3 wild-type and mutant constructs, images of 10 oocytes in each of 3 biological replicates (*n*=30 oocytes in total) were collected for analysis using a 20× air objective with a 1.2× digital zoom. Working from the top, left corner of the imaging dish, the first 10 oocytes for which *LE* localization was observed in the perinuclear cup, in L-bodies, and at the vegetal cortex were selected. Colocalization analysis was completed in ImageJ using the Colocalization threshold plugin using an ROI surrounding the oocyte.

### RNA immunoprecipitations (RIPs)

For the endogenous RIPs, approximately 600 stage II-III oocytes per protein condition were microinjected with 2 nL of 500 nM Vera-mCh, mCh-PTBP3 WT, or mCh-PTBP3 mut1234 RNA. Oocytes were cultured for 48 hours at 18°C in OCM+. Oocytes were then crosslinked with 0.1% formaldehyde in PBS for 10 min. at room temperature and then quenched for 5 min. in 250 mM glycine in 25 mM Tris (pH 7.4). Oocytes were then lysed in RIP buffer and clarified twice by centrifugation at 10,000×*g* for 10 min. at 4°C. 5 mg of antibody [anti-mCh (Abcam, ab62341) or Normal rabbit IgG (Sigma, NI01)] was added to each reaction and incubated for 1 hour at 4°C with rotation. Next, 15 µL of Pierce Protein A/G magnetic beads (#88803) in RIP buffer were added to each reaction and incubated for an additional 4 hours at 4°C, the beads were washed 3 times in RIP buffer, and bound proteins and RNAs were eluted from the beads in Pierce IgG Elution Buffer (ThermoFisher) via shaking (1200 rpm) at 24°C for 20 min. After removal of the eluent from the beads, crosslinking was reversed by incubation at 70°C for 45 min. To control for RNA extraction efficiency, 2.5 pg of luciferase RNA (Promega) was added to each sample prior to RNA extraction using RNeasy Plus Micro Kit (Qiagen). cDNA synthesis was performed using iScript cDNA synthesis kit (Biorad), and qRT-PCR was performed for *luciferase* and *vg1* RNAs using SybrGreen Powerup Master mix (ThermoFisher, A25742) per the manufacturer’s protocol using primers shown in Table S3.

For the barcoded RNA RIPs, 600 stage II-III oocytes per protein expressed were microinjected with 2 nL of 500 nM mCh-PTBP3 WT or mCh-PTBP3 mut1234 RNA and 150 nM *Barcode A-vg1 LE*, 150 nM *Barcode B-mut PTB LE*, and 150 nM *Barcode C-XBM* RNA. Oocytes were cultured and RIPs were performed as described above without crosslinking and decrosslinking of the samples. qRT-PCR was performed for *luciferase, Barcode A, Barcode B*, and *Barcode C* using primers shown in Table S3.

For the combinatorial PTBP3 RRM mutant RIPs, 200 stage II-III oocytes per protein expressed condition were microinjected with 2 nL of 500 nM mCh-PTBP3 WT, mCh-PTBP3 mut12, mCh-PTBP3 mut34, mCh-PTBP3 mut 3, mCh-PTBP3 mut 4, or mCh-PTBP3 mut1234 RNA and 150 nM *Barcode A-vg1 LE*. Oocytes were cultured and RIPs were completed as described above without crosslinking and decrosslinking of the samples. qRT-PCR was performed for *luciferase* and *Barcode A* using primers shown in Table S3.

### Recombinant protein expression and purification

MBP-PTBP3 fusion protein was expressed in BL21(DE3) *E. coli* (New England Biolabs) transformed with pET:THMT:PTBP3 (empty plasmid from Peti and Page, 2007) at 15°C overnight with 400 μM IPTG (Sigma). Induced pellets were resuspended in low imidazole purification buffer [20 mM sodium phosphate (pH 7.4), 1 M NaCl, 10 mM imidazole, and 1× Halt protease inhibitors (ThermoFisher)] and lysed via sonication. The suspension was cleared by centrifugation at 28,960 RCF for 30 min. at 4°C, and the resulting supernatant was bound to Ni-NTA resin (Qiagen) for 1 hour at 4°C, washed in 5 bed volumes of low imidazole purification buffer, and eluted in high imidazole purification buffer [20 mM sodium phosphate (pH 7.4), 1 M NaCl, 300 mM imidazole, and 1× Halt protease inhibitors (Thermo)]. Fractions containing MBP-PTBP3 were pooled, concentrated, buffer exchanged into storage buffer [20 mM sodium phosphate (pH 7.4), 250 mM NaCl], and flash frozen.

### In vitro *phase separation*

Purified MBP-PTBP3 protein was incubated with TEV protease for 2 hours at room temperature to cleave off the MBP tag. Protein for fluorescent imaging was labelled with AF647-NHS Ester (ThermoFisher) by resuspending the AF-647 in DMSO and incubating with cleaved PTBP3 protein for 1 hour at room temperature. Excess AF-647 was removed in a G50 desalting column (GE Healthcare). Labelled protein was buffer exchanged into 2× PTBP3 phase buffer [100 mM Tris (pH 7.5), 200 mM NaCl, 2 mM DTT]. Cleaved PTBP3 protein was diluted to 25 μM in 2× PTBP3 phase buffer with 10% fluorescent labelling for microscopy assays. 30 ng/μL stocks of *in vitro* transcribed RNA with 25% AF488-UTP labelled RNA in DEPC H_2_O for microscopy assays were denatured at 72°C for 10 min. and stored on ice. In a 20 μL reaction, 10 μL of PTBP3 protein (12.5 μM final protein concentration) and 10 μL of RNA (15 ng/μL final RNA concentration) were mixed and incubated for 1 hour at room temperature.

### Condensate morphology analysis

Condensates were imaged by placing 15 μL of the phase separation reaction onto a #1.5 coverslip with imaging spacers (Grace Bio-labs) and sealed with a slide (total imaging depth of ∼0.24 mM). Condensates were imaged on an inverted Olympus FV3000 confocal microscope using a 60× UPlan Super Apochromat objective (silicon oil, NA=1.3) and 5× digital zoom.

Condensate morphology was analyzed on 5 fields of view per replicate (*n*=15 fields of view total per RNA). Condensate number, size, and circularity were calculated based on manual thresholding of AF-647 labelled PTBP3 fluorescence using the “Analyze Particles” plugin in ImageJ. Condensates of any circularity and greater than 5 pixels in area were analyzed, excluding all condensates on the edge of the image. Partition coefficients (MCC) were calculated from the same analysis images using the “Colocalization Threshold” plugin in ImageJ.

### Turbidity Assays

For the NaCl concentration series, *in vitro* phase separation reactions were prepared as described above in 50 mM Tris (pH 7.5), 1 mM DTT, and the indicated NaCl concentration (25 mM, 100 mM, 250 mM, 500 mM, or 1 M). 12.5 μM PTBP3 was incubated with 0 ng/μL RNA (PTBP3 alone) or 15 ng/μL *LE* RNA (PTBP3 & *vg1 LE*). For the RNA concentration series, *in vitro* phase separation reactions were prepared as described above in 50 mM Tris (pH 7.5), 100 mM NaCl, 1 mM DTT. 12.5 μM PTBP3 (PTBP3 & *LE*) or buffer controls (*LE* alone,) were incubated with the indicated RNA concentration of *LE* RNA (0 ng/μL, 15 ng/μL, 37.5 ng/μL, and 75 ng/μL). Turbidity was assayed in a 384-well glass bottom plate (MatTek Corporation) with 20 μL samples sealed with clear optical film (Excel Scientific) to prevent evaporation. Absorbance of the samples at 600 nm was read using a Cytation 5 Multi-Mode Reader (BioTek) after 1 hour of incubation at room temperature. Absorbance data was normalized to buffer and TEV protease controls. In the turbidity assays, no fluorescent labels were used for either the protein or RNA.

### Immunoblotting

25 oocytes per mCh-PTBP3 construct and uninjected controls were homogenized in 50 uL of RIP buffer [25 mM Tris (pH 7.4), 0.5% NP40, 0.5 mM DTT, 150 mM KCl, 5 mM EDTA, 10 mM C_4_H_6_MgO_4_, 1× Halt protease inhibitors (ThermoFisher), and 2 nU/mL Ribolock RNase inhibitor (ThermoFisher)]. Lysates were clarified via centrifugation at 10,000×*g* for 10 min. at 4°C and then boiled in Laemmli sample buffer (Laemmli, 1970). For immunoblotting, anti-mCh (Abcam, ab62341) and anti-tubulin (Abcam, ab4074) primary antibodies were used at 1:1000. Secondary goat anti-rabbit IgG-HRP antibody (Abcam, 97200) was used at 1:15,000. Blots were developed using SuperSignal West Pico PLUS chemiluminescent substrate (ThermoFisher) and imaged using a Biorad ChemiDoc.

### Fluorescence recovery after photobleaching (FRAP)

For the oocyte FRAP, Stage II oocytes were microinjected with 2 nL of 500 nM RNA encoding mCh-PTBP3 and 250 nM Cy5-UTP labelled *LE* RNA for FRAP of PTBP3 wild-type and mutant proteins. For RNA FRAP, 2 nL of 250 nM Cy3-UTP labelled test RNA (*LE, mut PTB LE*, or *XBM*) was microinjected into stage II oocytes, along with 250 nM Cy5-UTP labelled *LE* RNA to mark the L-bodies. Microinjected oocytes were cultured for 48 hours in OCM+. 7 oocytes per biological replicate (*n*=21 oocytes total per construct tested) were analyzed. A 10 μm^2^ ROI was bleached using the 488 nm laser at 100% for 2 seconds. Fluorescence recovery was monitored every 5 seconds for 100 iterations. FRAP calculations were completed as previously described (Gagnon et al., 2013) and data was analyzed via one phase non-linear regression using GraphPad Prism 9.

For the *in vitro* condensate FRAP, 7 condensates per replicate (*n*=21 condensates total per RNA) were analyzed. A 2 μm^2^ ROI was bleached using the 405 nm laser at 50% and the 561 nm laser at 100% for 0.8 seconds. Fluorescence recovery was monitored every 5 seconds for 50 iterations. FRAP calculations were completed as previously described (Gagnon et al., 2013) and data was analyzed via one phase non-linear regression using GraphPad Prism 9.

## Acknowledgements

We thank Nicolas L. Fawzi, Erica N. Larschan, Jessica P. Otis, and Liam C. O’Connell for comments on the manuscript. Thank you to Nicolas L. Fawzi for gifting the *in vitro* protein expression plasmid and advice on recombinant protein expression and purification. This work was funded by R01GM071049 from the NIH to KLM. The authors declare no competing financial interests.

## Author Contributions

SEC and KLM designed, carried out, and analyzed experiments. SEC and KLM drafted and revised the manuscript.

## Supplemental Figure Legends

**Figure S1:**
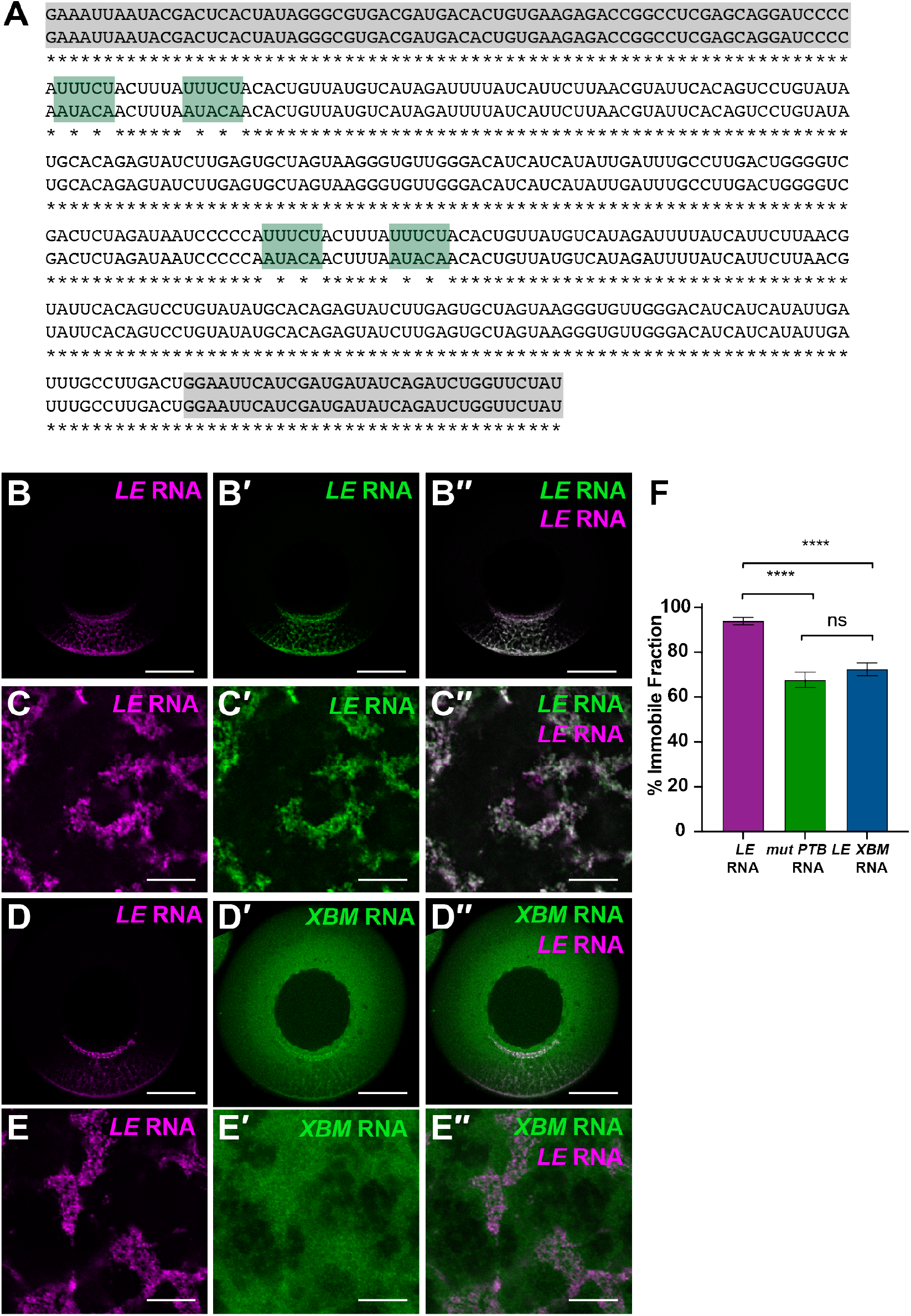
Comparison of localized and non-localized RNAs in L-body enrichment and dynamics. Related to Fig. 1 **(A)** Alignment of *LE* RNA (top) and *mut PTB LE* RNA (bottom) with identical nucleotides indicated by *. PTB binding sites highlighted in green and nonspecific sequences added for length matching and *in vitro* transcription are shown in grey. **(B)** Whole mount fluorescence of a stage II oocyte microinjected with Cy-5-labeled *LE* RNA (A, magenta) and Cy3-labeled *LE* RNA (A’, green). Overlap is shown in A’’; scale bars=100 µm. **(C)** High magnification view of L-bodies in a stage II oocyte microinjected with Cy-5-labeled *LE* RNA (C, magenta) and Cy3-labeled *LE* RNA (C’, green). Overlap is shown in C’’; scale bars=10 µm. **(D)** Whole mount fluorescence of a stage II oocyte microinjected with Cy-5-labeled *LE* RNA (D, magenta) and Cy3-labeled *XBM* RNA (D’, green). Overlap is shown in D’’; scale bars=100 µm. **(E)** High magnification view of L-bodies in a stage II oocyte microinjected with Cy-5-labeled *LE* RNA (E, magenta) and Cy3-labeled *XBM* RNA (E’, green). Overlap is shown in E’’; scale bars=10 µm. **(F)** Percent immobile fraction for RNA FRAP experiments shown in (Figure 1E). Error bars represent standard error of the mean. ns indicates p> 0.5, **** indicates p<0.001.

**Figure S2:**
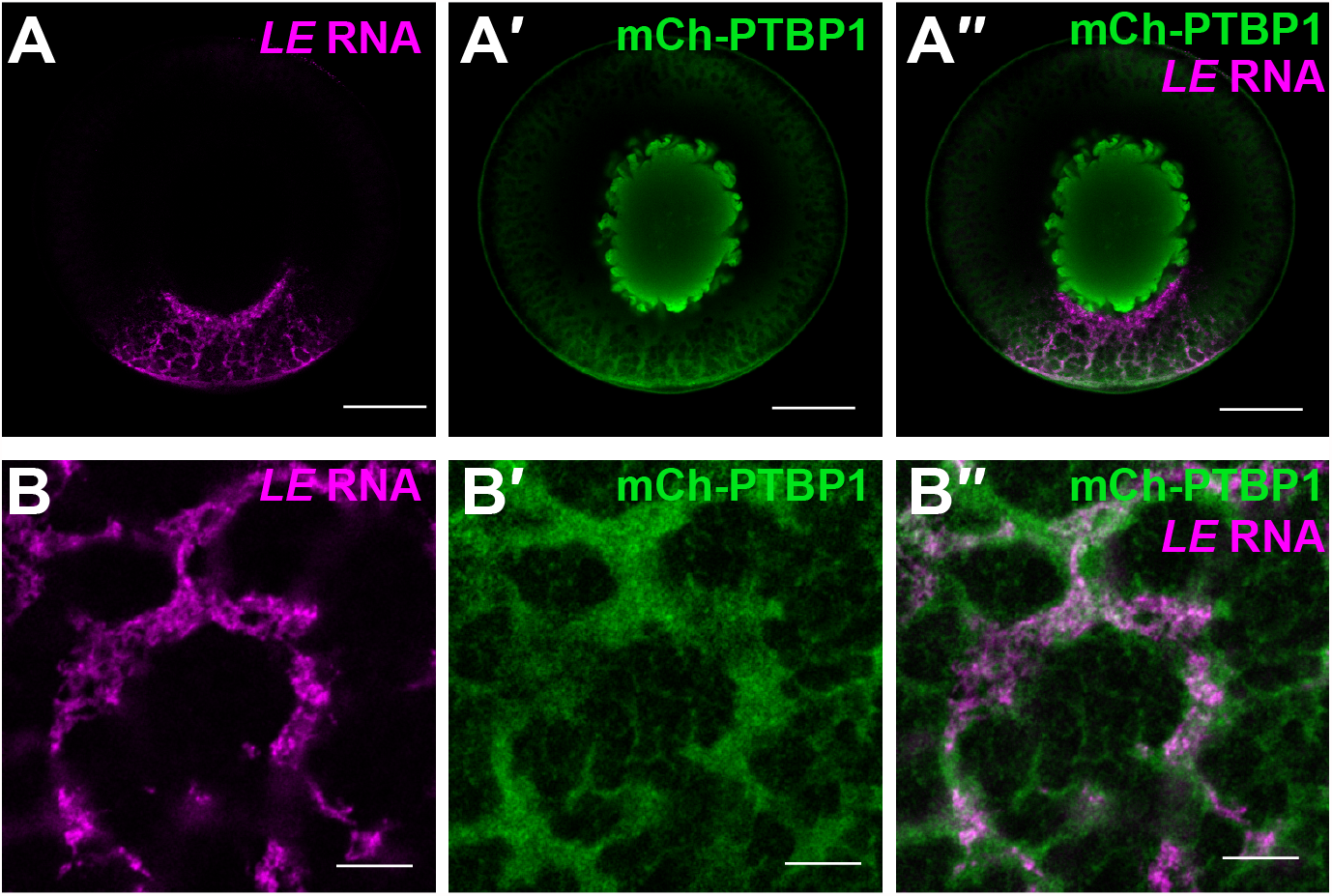
PTBP1 is a nuclear and cytoplasmic paralog of PTBP3. Related to Fig. 2 **(A)** Fluorescently-labeled *LE* RNA (A, magenta), was microinjected into a stage III oocyte expressing and mCh-PTBP1 (A’, green), as detected by anti-mCh IF. Overlap is shown in A’’. The vegetal pole is at the bottom and the scale bars=100 µm. **(B)** High magnification view of L-bodies in a stage III oocyte microinjected with *LE* RNA (B, magenta) and expressing mCh-PTBP1, as detected by anti-mCh IF (B’, green). Overlap is shown in B’’; scale bars=10 µm.

**Figure S3:**
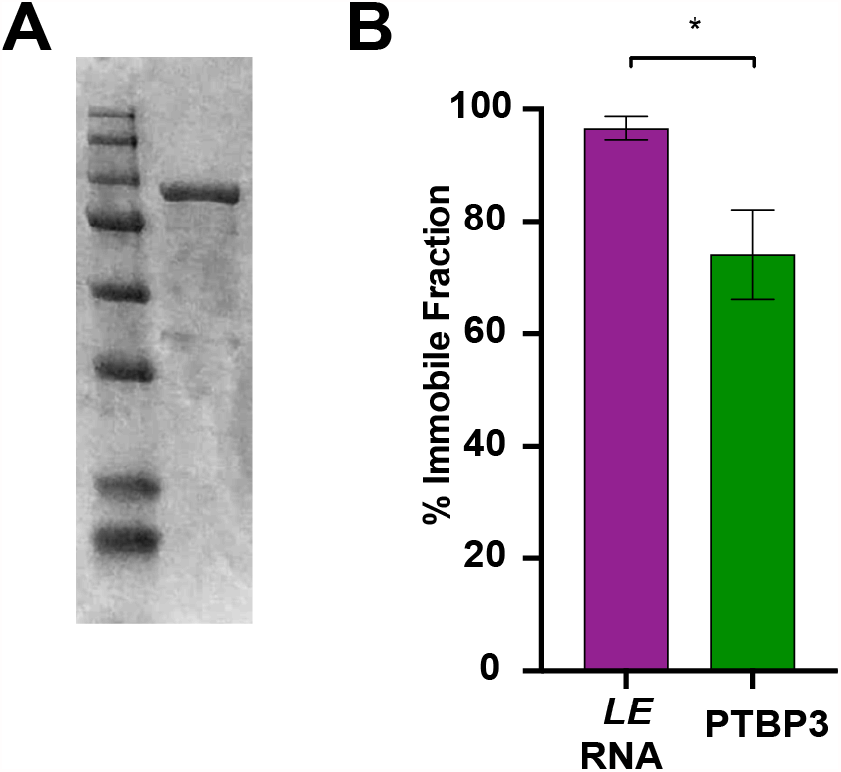
*In vitro* phase separation with recombinant PTBP3. Related to Fig. 3. **(A)** Coomassie stained gel showing recombinant MBP-PTBP3 (102 kDa) purification. **(B)** Percent immobile fraction analysis for FRAP shown in (Fig. 3D). Error bars represent standard error of the mean. * indicates p<0.5

**Figure S4:**
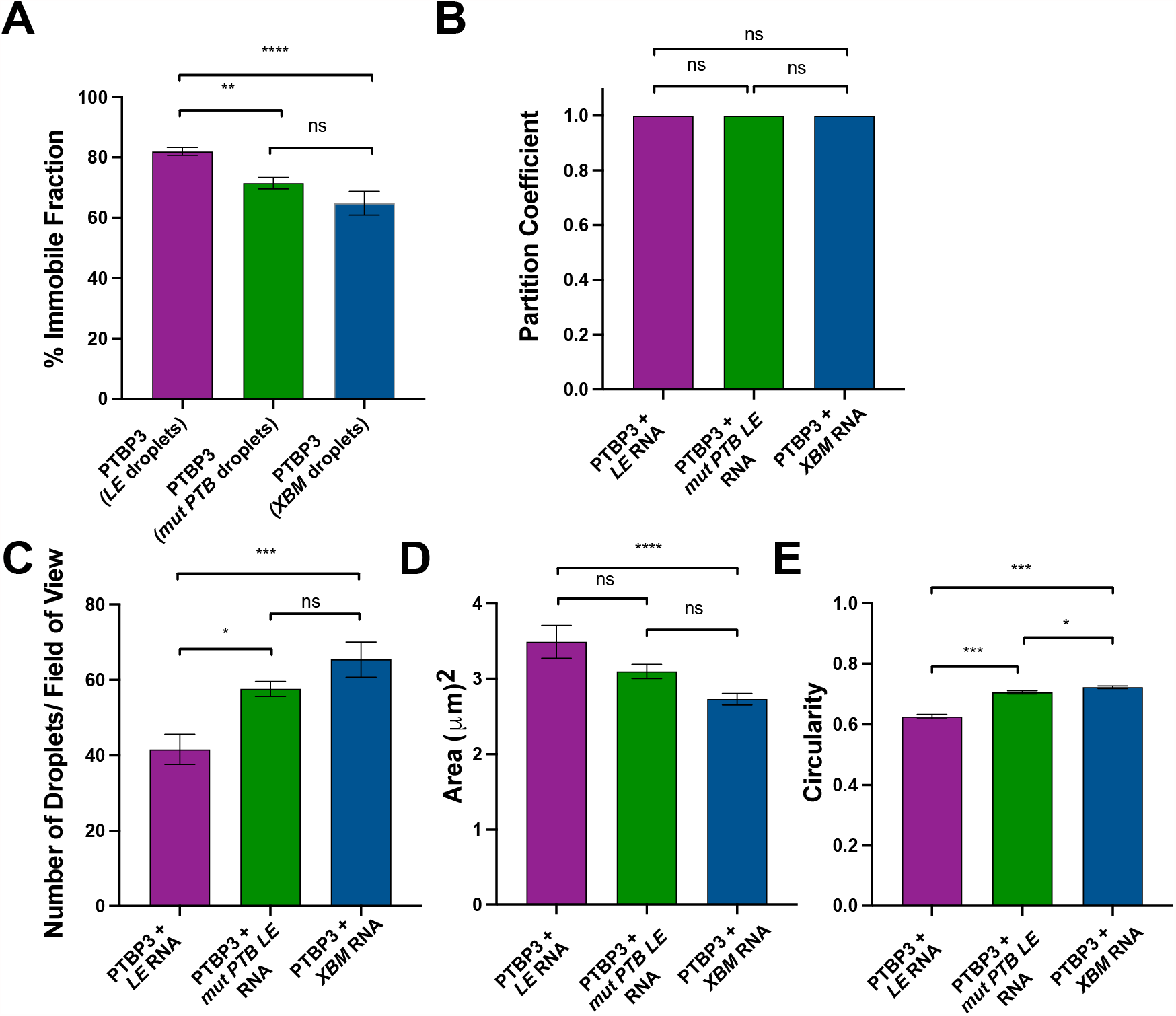
Quantification of *in vitro* PTBP3 condensates with *LE, mut PTB LE*, or *XBM* RNAs. Related to Fig. 4 **(A)** Percent immobile fraction for FRAP analyses shown in (Fig. 4E). Error bars represent standard error of the mean. * indicates p<0.5 **(B)** Partition coefficients for PTBP3 with *LE* RNA (magenta), *mut PTB LE* RNA (green), and *XBM* RNA (blue). Error bars represent standard error of the mean. *n*=15 fields of view, ns indicates p>0.05. **(C)** The average number of condensates per field of view for PTBP3-*LE* RNA (magenta), PTBP3-*mut PTB LE* RNA (green), and PTBP3-*XBM* RNA (blue) was determined by analysis of images such as shown in (Fig. 4A-C). Error bars represent standard area of the mean, *n*=15 fields of view. ns indicates p>0.5, * indicates p<0.5, *** indicates p<0.01. **(D)** The average area of condensates formed by PTBP3-*LE* RNA (magenta), PTBP3-*mut PTB LE* RNA (green), and PTBP3-*XBM* (blue) was determined by analysis of images such as shown in (Fig. 4A-C). Error bars represent standard area of the mean, *n*=15 fields of view. ns indicates p>0.5, **** indicates p<0.001. **(E)** The average circularity of condensates formed by PTBP3-*LE* RNA (magenta), PTBP3-*mut PTB LE* (green), and PTBP3-*XBM* (blue) was determined by analysis of images such as shown in (Fig. 4A-C). Error bars represent standard area of the mean, *n*=15 fields of view. * indicates p<0.5, *** indicates p<0.01.

**Figure S5:**
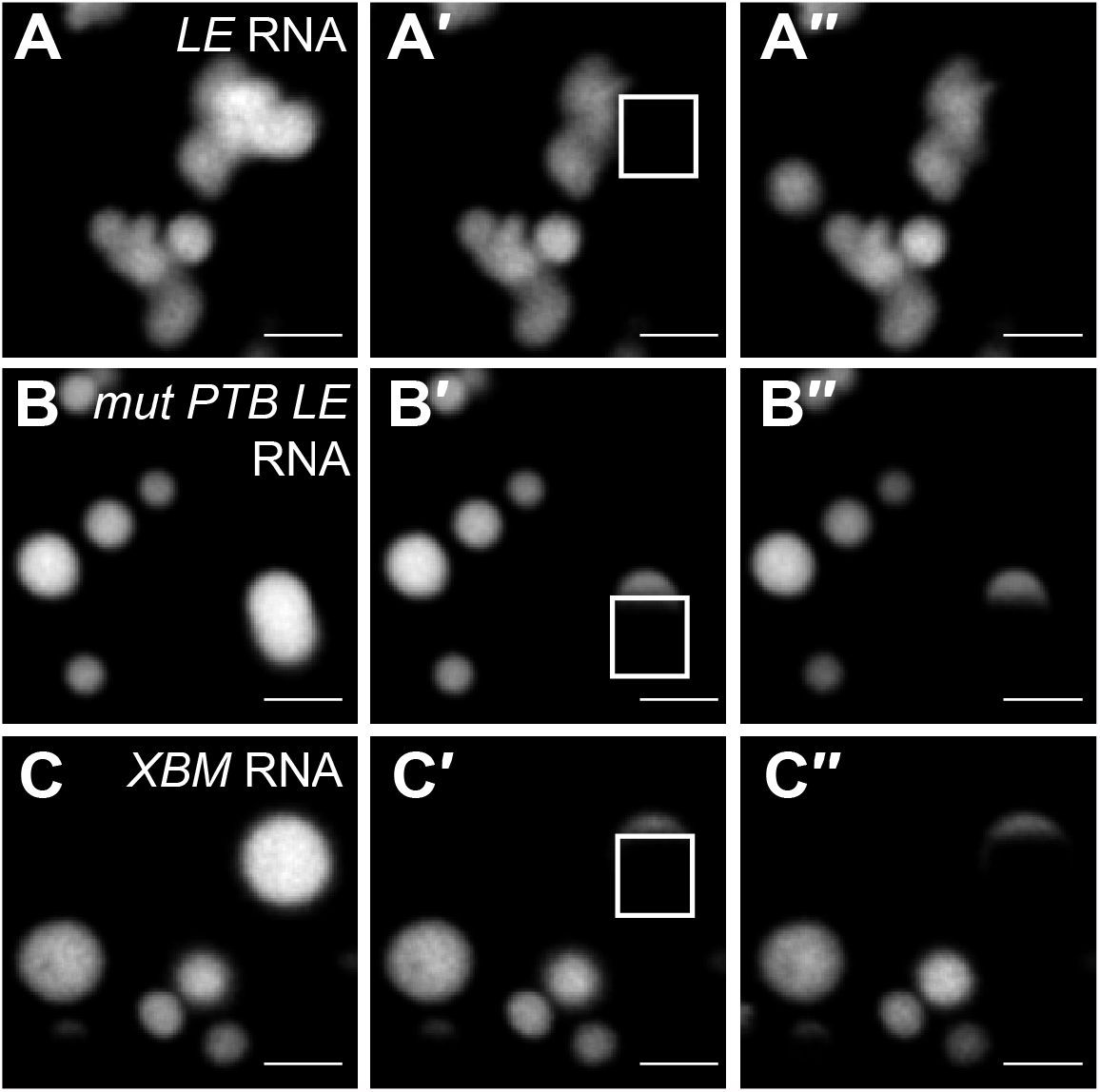
Partial condensate FRAP shows that RNAs in *in vitro* condensates are non-dynamic. Related to Fig. 4. **(A)** An image of *LE* RNA in PTBP3-*LE* condensates is shown with a 2 µm^2^ FRAP ROI (white) over part of the condensate. (A) *LE* RNA fluorescence pre-bleach, post-bleach (A’), and 4 min. post-recovery (A’’). Scale bars=2 µm **(B)** An image of *mut PTB LE* RNA in PTBP3-*mut PTB LE* condensates is shown with a 2 µm^2^ FRAP ROI (white) over part of the condensate. (B) *mut PTB LE* RNA fluorescence pre-bleach, post-bleach (B’), and 4 min. post-recovery (B’’). Scale bars=2 µm **(C)** An image of *XBM* RNA in PTBP3-*XBM* condensates is shown with a 2 µm^2^ FRAP ROI (white) over part of the condensate. (C) *XBM* RNA fluorescence pre-bleach, post-bleach (C’), and 4 min. post-recovery (C’’). Scale bars=2 µm

**Figure S6:**
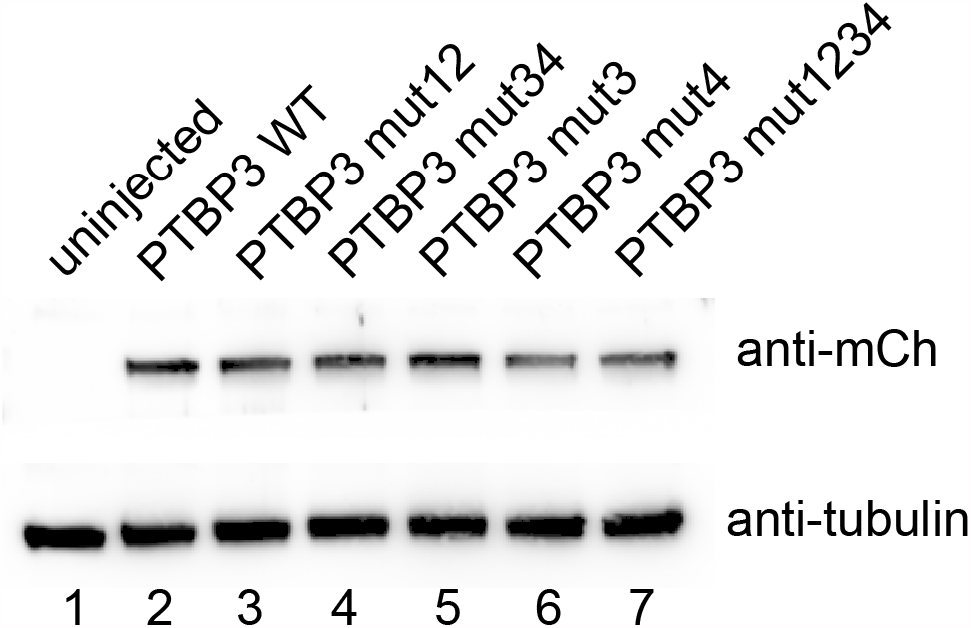
Expression of PTBP3 RRM mutants in oocytes. Related to Fig. 5. Immunoblot analysis was carried out using lysates prepared from stage II oocytes injected with mCh-PTBP3 WT or RRM mutants after centrifugation to assay solubility. Anti-mCh (top) and anti-tubulin (bottom) blots are shown. Proteins expressed are as follows: Lane 1: uninjected control, Lane 2: mCh-PTBP3 WT, Lane 3: mCh-PTBP3 mut12, Lane 4: mCh-PTBP3 mut34, Lane 5: mCh-PTBP3 mut3, Lane 6: mCh-PTBP3 mut4, Lane 7: mCh-PTBP3 mut1234.

## Supplemental Tables

**Table S1:**
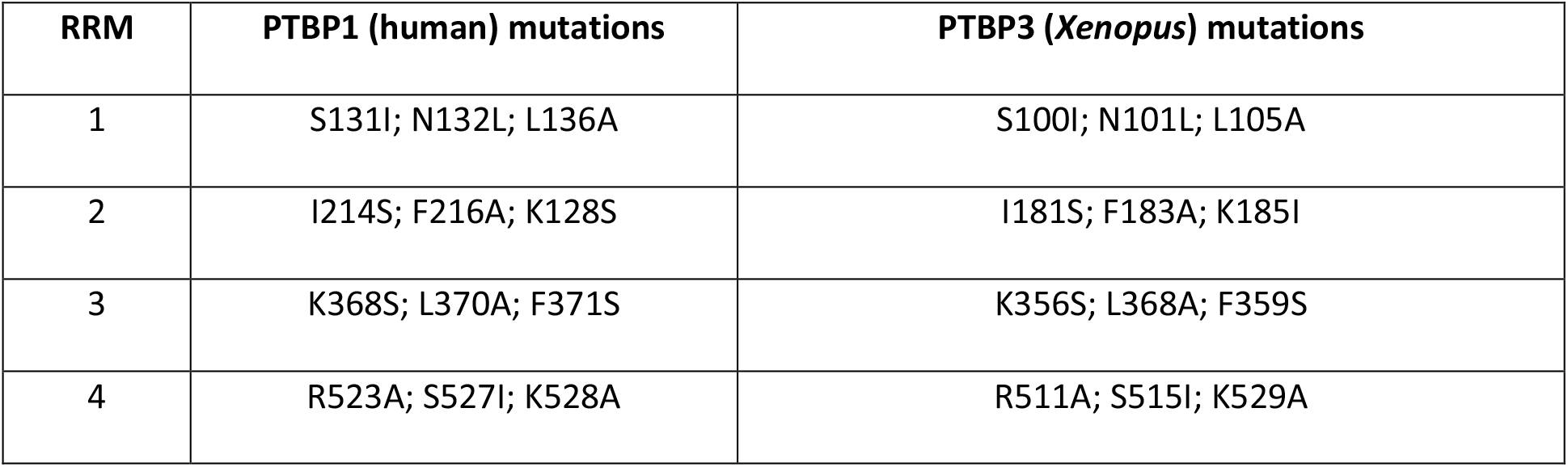
List of protein mutations introduced to PTBP3 for each of the four RRMs. Related to Fig. 2 and Fig.5.

**Table S2:**
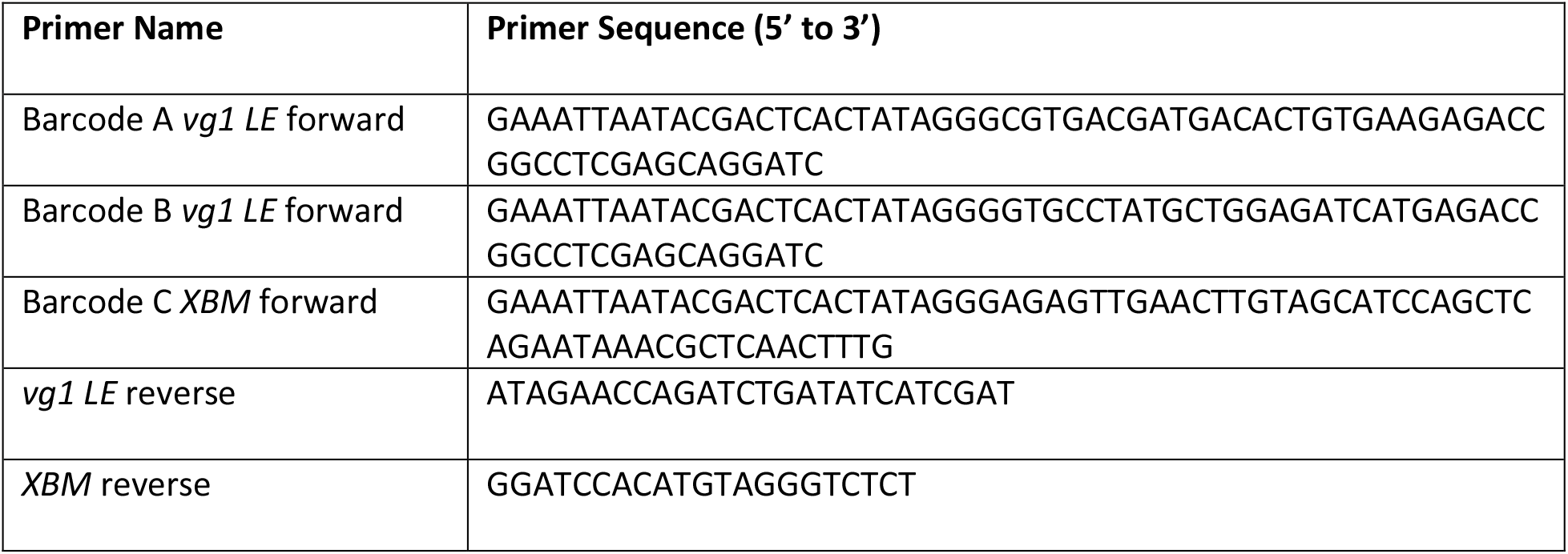
PCR primers. Related to methods.

**Table S3:**
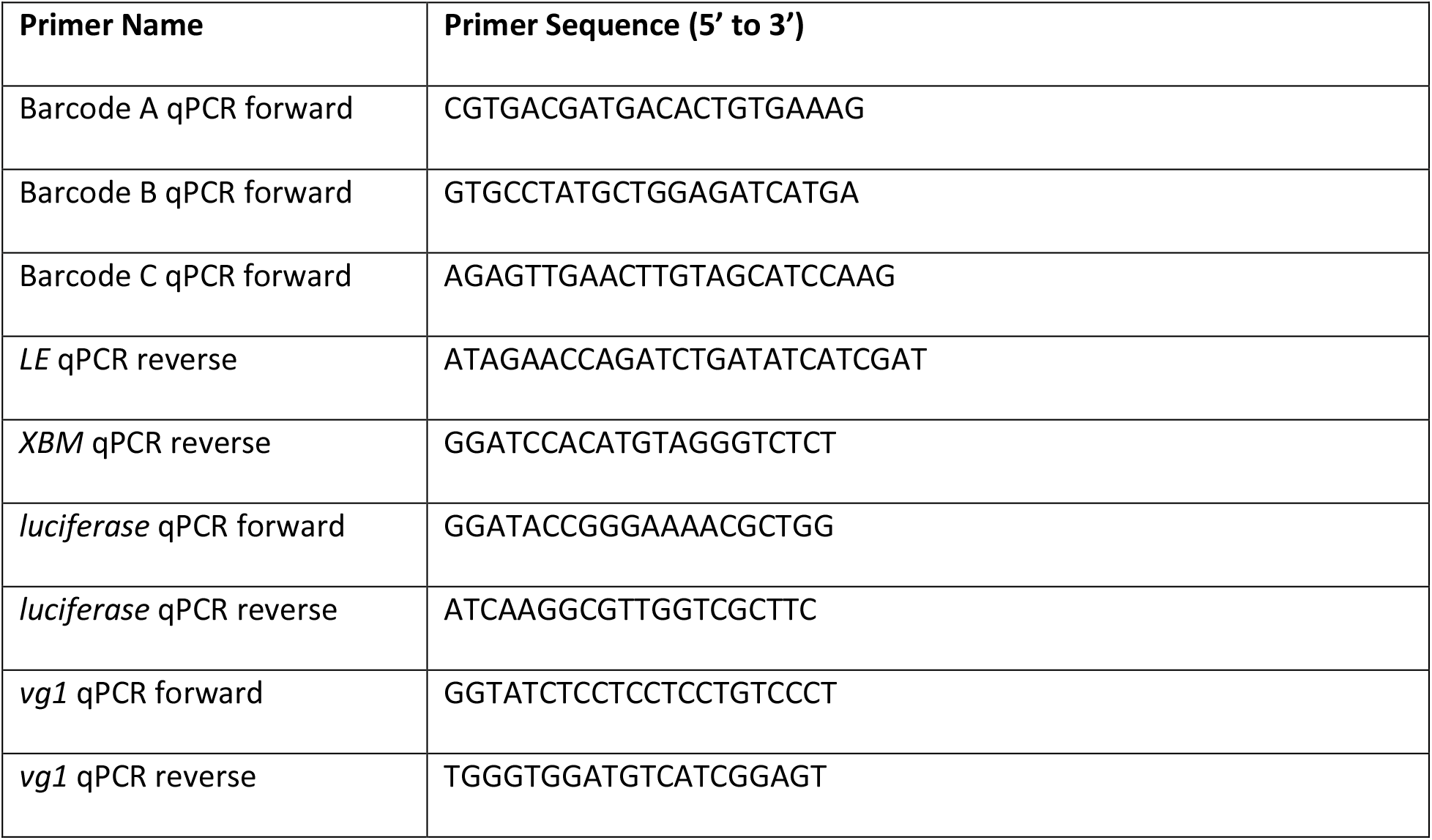
qPCR primers. Related to methods.

